# A CRISPR-based genome-wide screen for adipogenesis reveals new insights into mitotic expansion and lipogenesis

**DOI:** 10.1101/2020.07.13.201038

**Authors:** Rachel E. Turn, Keren I. Hilgendorf, Carl T. Johnson, Kyuho Han, Atefeh Rabiee, Janos Demeter, Mohammad Ovais Aziz-Zanjani, Pablo Domizi, Ran Cheng, Yingdi Zhu, Zewen Jiang, Katrin J. Svensson, Michael C. Bassik, Peter K. Jackson

**Affiliations:** Baxter Laboratory, Department of Microbiology & Immunology, Stanford University School of Medicine, Stanford, CA 94305, USA; Department of Medicine, Stem Cell and Regenerative Medicine Program, Stanford University School of Medicine, Stanford, CA 94305, USA; Department of Genetics, Stanford University School of Medicine, Stanford, CA 94305, USA; Department of Chemical and Systems Biology, Stanford University, Stanford, CA 94305, USA; Department of Pediatrics, Stanford University School of Medicine, Stanford, CA 94305, USA; Department of Biology, Stanford University, Stanford, CA 94305, USA; Laboratory of Physical and Analytical Electrochemistry (LEPA), Ecole Polytechnique Fédérale de Lausanne (EPFL) Valais Wallis, Rue de l’Industrie 17, 1950 Sion, Switzerland; Department of Pathology, Stanford University, Stanford, CA 94305, USA; Stanford Diabetes Research Center, Stanford University School of Medicine, Stanford, CA, 94305, USA

**Keywords:** Adipogenesis, obesity, mitotic expansion, lipogenesis, CRISPR, ubiquitin, lipid, hypusine, Nedd8, translation

## Abstract

In response to excess nutrients, white adipose tissue expands by both generating new adipocytes and by upregulating lipogenesis in existing adipocytes. Here, we performed a genome-wide functional genomics screen to identify regulators of adipogenesis in the mouse 3T3-L1 preadipocyte model. The pooled screening strategy utilized FACS to isolate populations based on lipid content by gating for fluorescence intensity of the lipophilic, green fluorescent BODIPY dye. Additionally, the approach categorized if genes functioned during mitotic expansion or lipogenesis. Cellular mechanisms regulating the rate of protein translation and protein stability were found critical for adipogenesis and lipogenesis. These protein-directed mechanisms were further supported by proteomic analyses, which demonstrated that essential changes in protein abundance driving 3T3-L1 adipogenesis were not driven by transcription. We exemplify this theme by showing that the hypusination pathway, a conserved regulator of translation initiation, is critical to translate adipogenic inducers of mitotic expansion and that the neddylation/ubiquitin pathway modulates insulin sensitivity to regulate lipogenesis.

## Introduction

Obesity, characterized by an excessive expansion of white adipose tissue, is a world-wide epidemic; its comorbidities, including diabetes, cancer, cardiovascular and hepatic disease, pose a significant and costly challenge to public health (Bluher, 2019). White adipose tissue expands in response to overnutrition by both increasing the size of existing adipocytes and by generating new adipocytes via *de novo* adipogenesis of preadipocytes (Haczeyni et al., 2018). Notably, white adipose tissue which consists of smaller adipocytes, even if greater in number, is metabolically healthier as it reduces the risk of inflammation and downstream complications (Ghaben and Scherer, 2019). Thus, there is substantial interest in understanding the molecular mechanisms that selectively regulate mitotic expansion versus lipogenesis.

The process of adipogenesis can be functionally subdivided into several distinct processes: 1) activation of preadipocytes, 2) expansion of preadipocyte numbers, 3) differentiation into an adipocyte, and 4) lipogenesis (Rosen et al., 2000; Tang et al., 2003b). Much of our current understanding of adipogenesis has been gained from the study of a murine cell line adipogenesis model, 3T3-L1 cells (Green and Meuth, 1974). These immortalized cells differentiate robustly and uniformly upon addition of a differentiation cocktail, greatly enabling biochemical and genetic characterization of adipogenesis. Specifically, in response to the differentiation factors insulin, dexamethasone, and 3-isobutyl-1-methylxanthine (IBMX), confluency-arrested 3T3-L1 cells re-enter the cell cycle, undergo ∼2 rounds of specialized mitosis (termed mitotic clonal expansion, MCE), and then activate a well-characterized transcriptional cascade before accumulating lipids in response to insulin (Zebisch et al., 2012). A multitude of transcription factors and transcriptional coregulators of adipogenesis have been identified, which in turn activate the master adipogenic transcription factors C/EBPα and PPARγ (MacDougald and Lane, 1995). After induction with insulin and cAMP signaling, non-transcriptional mechanisms regulating steps in adipogenesis have received less attention.

*In vivo*, quiescent preadipocytes are activated in response to physiological cues such as dietary fatty acids (FAs) and insulin to re-enter the cell cycle to self-renew and to differentiate into adipocytes (Hilgendorf et al., 2019; Jeffery et al., 2015). In contrast to this asymmetric cell division, 3T3-L1 MCE is symmetric with both daughter cells uniformly differentiating into adipocytes, an advantage for both biochemical and genetic analyses. Though the use of 3T3-L1 immortalized preadipocytes may not reflect precisely all of the variables impinging on adipogenesis in living fat tissue, 3T3-L1 to date serve as a standard for probing the mechanisms driving adipogenesis as many of the pathways discovered in 3T3-L1 recapitulate in cultured primary preadipocytes. Of note, while some other cell line models of adipogenesis appear to not undergo complete mitoses, these cells still need to progress through S-phase (Cho and Jefcoate, 2004). We here reaffirm the importance of cell cycle regulation in multiple models of adipogenesis including primary human preadipocytes. Thus, the replication of DNA, both *in vivo* and *in vitro,* is a critical though poorly understood stage of adipogenesis. Lipogenesis is a well-regulated metabolic process that converts excess energy into triglycerides, the predominant constituent of lipid droplets in white adipocytes. This process is required for both the formation of lipid droplets in newly differentiating preadipocytes as well as for storing additional energy in existing adipocytes, increasing their size. In response to insulin, the differentiating preadipocyte/expanding adipocyte takes up excess energy (-e. g. glucose, FAs) and activates enzymatic pathways leading to the formation of acetyl-CoA and malonyl-CoA, the building blocks of FAs (Song et al., 2018). Esterification of FAs to glycerol leads to the formation of triglycerides to form or grow lipid droplets. Since lipid droplets are highly dynamic, composed of a multitude of macromolecules, and are critical to protect cells from lipotoxicity, their biology has become the focus of intense investigation (Olzmann and Carvalho, 2019; Walther and Farese, 2012).

We sought to systematically identify genes involved in adipogenesis by conducting a genome-wide screen using CRISPR/Cas9 to disrupt gene function. To date, it has been difficult to characterize the full array of adipogenic genes due to the technical limitations of performing large scale screens using arrayed libraries (Guo et al., 2008; Sohle et al., 2012). Alternatively, pooled screening formats require a mechanism to isolate cell populations with a desired phenotype. Consequently, most studies to date have used pooled screening formats to screen for genes that directly affect cell growth or cell death or have used engineered reporter cell lines to establish a selection platform to screen for genes of interest indirectly (van Beekum et al., 2012).

Here, we have developed a lipid-based sorting strategy to directly isolate differentiated adipocytes versus undifferentiated preadipocytes. We used 3T3-L1 preadipocytes expressing Cas9 constitutively and a single guide RNA (sgRNA) lentiviral library targeting the mouse genome. By differentiating the sgRNA-infected pool of 3T3-L1 cells and sorting based on lipid content, we identified novel adipogenic regulators and characterized their relative importance to mitotic clonal expansion versus lipogenesis. Intriguingly, we further show that cellular mechanisms regulating translation and protein stability are critical for adipogenic fate change. We illustrate the strength of this screen by describing how hypusination, a key regulator of translation initiation, and neddylation, central to many E3 ubiquitin ligases driving ubiquitin-dependent proteolysis, regulate MCE and lipogenesis, respectively.

## Results

### A lipid-based sorting strategy enables pooled screening for regulators of adipogenesis

Pooled CRISPR/Cas9 screening strategies are based on the ability to isolate cell populations which enrich or deplete for a desired phenotype, followed by deep sequencing to quantify changes in relative sgRNA abundance between populations. Since the ability to differentiate does not affect relative cell abundance in a population, we instead developed a sorting-based strategy based on fluorescent intensity of the lipophilic, green fluorescent dye BODIPY 493/503 (Fig. 1A). Briefly, 3T3-L1 preadipocytes expressing Cas9 protein were infected at low multiplicity of infection with a previously developed mouse sgRNA library (Breslow et al., 2018; Morgens et al., 2017). This library covers the entire mouse genome with 10 sgRNAs per gene as well as more than 10,000 non-targeting/safe sgRNAs. To avoid a bottleneck effect, the 3T3-L1 pool was maintained at ∼1000x coverage per sgRNA at all times (∼200 million cells in total). The 3T3-L1 pool was passaged for more than 2 weeks prior to initiating the experiment to select against sgRNAs targeting genes essential for viability, allowing them to drop out of the pool. The surviving pool of 3T3-L1 cells (∼97% of initial sgRNA library) was grown to confluence and differentiated upon addition of differentiation cocktail. After 4 days of differentiation, the 3T3-L1 pool was stained with BODIPY followed by large scale cell sorting for green fluorescence by flow cytometry. The Day 4 time point was chosen, since adipocytes at this time point contain small lipid droplets to mark differentiation, but these are not so large as to hinder sorting.

**Figure 1:**
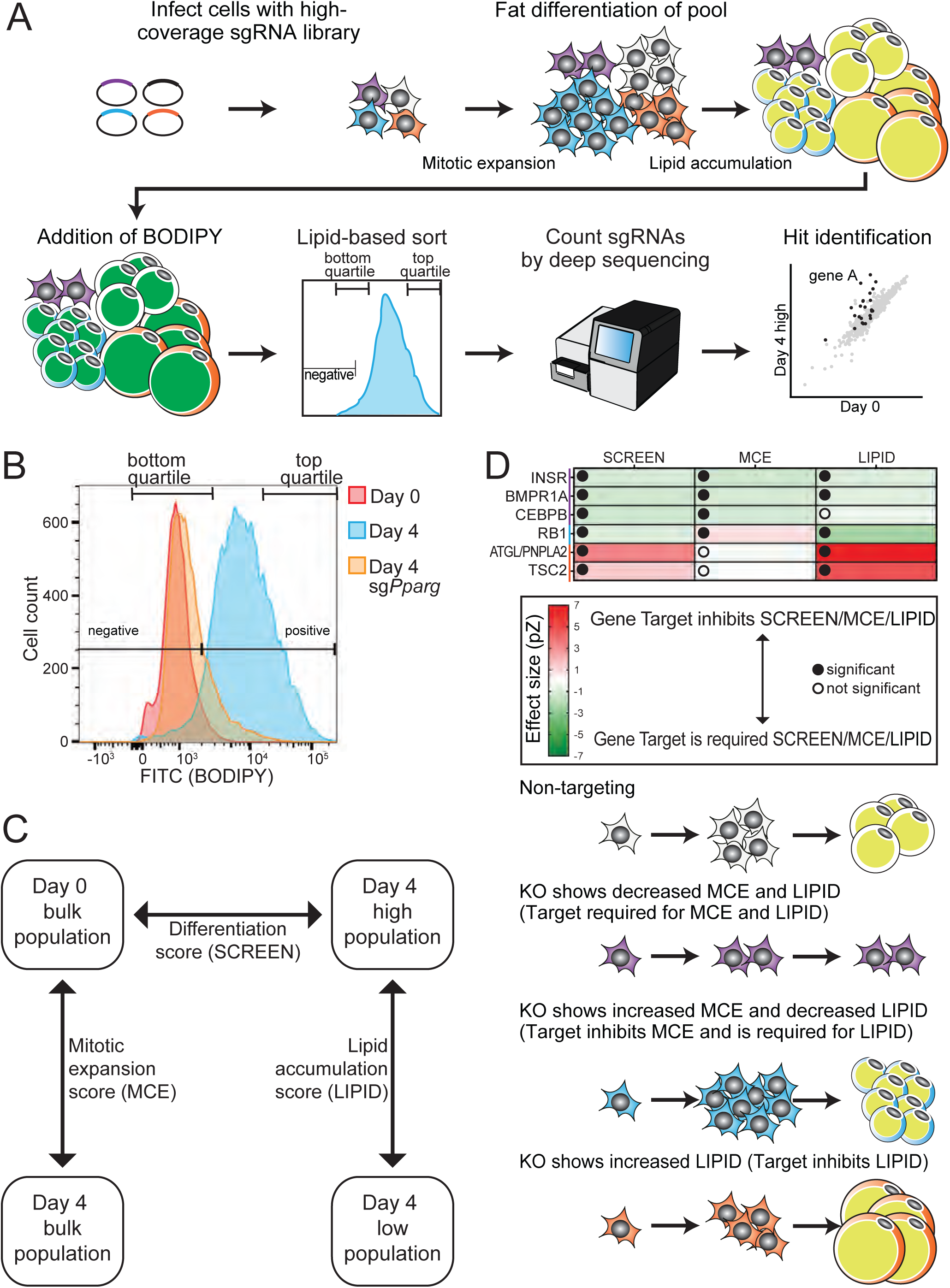
Development of an adipogenesis screen using a lipid-based sorting strategy. a. Overview of screening strategy. 3T3-L1 cells receiving a non-targeting control sgRNA are shaded in white, cells receiving positive or negative regulators of adipogenesis are shaded in color. Lipophilic green-fluorescence BODIPY dye is denoted in green. b. Flow cytometry histograms of undifferentiated and differentiated 3T3-L1 cells treated with BODIPY. Cells lacking PPARG fail to differentiate and remain BODIPY-negative. Top and bottom quartile were sorted to obtain a Day 4 low and Day 4 high population. c. Phenotypic scores were calculated by comparing sgRNA abundance relative to non-targeting control sgRNAs between two distinct cell populations. The differentiation score (SCREEN) is calculated by comparing Day 0 to Day 4 high, the MCE score is calculated by comparing Day 0 to bulk (unsorted) Day 4, and the lipid accumulation score (LIPID) is calculated by comparing Day 4 low to Day 4 high. d. Heatmap displays phenotypic scores obtained for known adipogenic regulators regulating MCE and/or LIPID. Schematic representation of results for each category. Cells receiving a non-targeting control sgRNA are shaded in white, cells lacking positive regulators of MCE are shaded in yellow, cells lacking a negative regulator of MCE which is also a positive regulator of LIPID are shaded in blue, and cells lacking a negative regulator of LIPID are shaded in red.

First, we confirmed the robustness and uniformity of adipogenesis as well as our ability to isolate differentiated adipocytes versus undifferentiated preadipocytes by FACS in parental 3T3-L1 cells. The staining protocol separated the two populations based on green fluorescent intensity with only ∼5% of cells falling into the incorrect category (negative and positive, Fig. 1B). As proof of principle, we infected 3T3-L1 Cas9 cells with a sgRNA targeting the master adipogenic transcription factor PPARγ. Consistent with PPARγ being required for adipogenesis, the bulk of these cells fell into the negative category even after 4 days of differentiation (Fig. 1B, yellow). Thus, this screening strategy is capable of isolating cell populations which enrich or deplete for the presence of intracellular lipid and therefore the ability to undergo differentiation.

Two sequential rounds of screening were performed in duplicates (Fig. S1A). First, we used the genome-wide sgRNA library and gated for the top 50% BODIPY positivity. This more lenient cutoff was chosen to enable sorting of a sufficiently large cell population to maintain a ∼1000x coverage per sgRNA even in the sorted cell population. Genes affecting adipogenesis were identified by quantifying and comparing sgRNA abundance in the Day 4 BODIPY^top^ ^50%^ cell population versus the bulk Day 0 cell pool. Effect size was calculated using the Mann-Whitney U test for the 10 sgRNAs per gene target compared to the non-targeting/safe sgRNA population, and the significance cut-off was set at an estimated FDR of 5% (Storey and Tibshirani, 2003). This first screen identified ∼1500 gene targets as candidate regulators of adipogenesis (Table S1). We selected 10 of these gene targets for manual confirmation. Selection was based on gene targets scoring as highly significant, but having little to no previously known function in regulating adipogenesis. We confirmed that CRISPR-mediated loss of 9 out of the 10 gene targets inhibited adipogenesis, including two gene targets whose loss had comparable effects to loss of C/EBPβ and PPARγ (Fig. S1B). As an additional confirmation, two high scoring yet unknown genes, E130309D02RIK and GM14393, were knocked out in 3T3-L1 cells, which subsequently failed to properly differentiate (Fig. S1C). This argues that the list of gene targets obtained from screen 1 is strongly enriched for regulators of adipogenesis.

A second round of screening was conducted under more stringent sorting conditions to identify strong and high-confidence regulators of adipogenesis. Briefly, we developed a new custom library targeting the top 1500 hits of the first screen as well as additional genes of interest related to signal transduction pathways (Fig. S1A). We repeated the screening strategy in duplicates, this time sorting for both the top and bottom quartile of BODIPY signal intensity. Additionally, bulk cell populations on day 0 and day 4 (unsorted) were collected. Together, these four cell populations allowed for the identification of three different but related properties that can contribute to overall BODIPY signal (Fig. 1C, Table S2). Genes affecting differentiation (henceforth referred to as SCREEN) were identified by comparing sgRNA abundance in the Day 4 BODIPY^high^ cell population versus the bulk Day 0 cell pool, reflecting depletion or enrichment of sgRNAs in the most clearly differentiated population. Genes affecting mitotic expansion (MCE) were identified by comparing sgRNA abundance in the bulk Day 4 versus the bulk Day 0 cell populations. Genes affecting lipid accumulation (henceforth referred to as LIPID) were identified by comparing sgRNA abundance in the Day 4 BODIPY^high^ versus BODIPY^low^ cell populations, allowing us to distinguish direct effects on lipolysis and lipogenesis irrespective of MCE. Taken together, the second screen not only identified putative regulators of adipogenesis but categorized their relative importance to MCE versus LIPID. Notably, the first screen successfully culled the number of adipogenic regulators found in the whole genome, because hits from the first screen were significantly enriched among the hits identified for SCREEN, MCE, and LIPID (Fig. S1D).

Since adipogenesis is a function of both the ability to undergo MCE and to accumulate lipid, we next sought to establish how the three phenotypic scores relate to each other. Interestingly, the size of the SCREEN effect strongly correlated with the simple sum of the MCE and LIPID effects (Fig. S1E, F). This argues that the ability to categorize adipogenic genes into regulators of MCE versus LIPID can provide significant functional discrimination for assigning the molecular mechanism of action for both known and novel hits identified in this screen. To validate this model, we investigated adipogenic genes known to regulate MCE versus LIPID. The insulin receptor, the BMP1A receptor, and C/EBPβ are known to be required for adipogenesis, and all three are known inducers of MCE (Bowers and Lane, 2007; Gregoire et al., 1998; Tang et al., 2003b). Consistently, all three genes showed significant, SCREEN and MCE negative scores or loss of function effects (Fig. 1D purple in heatmap and cartoon). In addition, the ligands insulin and BMP4 promote lipid accumulation (Kersten, 2001; Modica et al., 2016), and their associated receptors (INSR and BMPR1A receptors respectively) also showed a significant negative LIPID effect (Fig. 1D purple). Similarly, our screen validated that the cell cycle regulator RB1 is required for adipogenesis, but this was a consequence of dual, opposing roles for RB1, with: (1) a significant positive MCE effect, consistent with it being required to exit MCE; and (2) a significant negative LIPID effect, consistent with RB1 promoting expression of lipogenic genes (Fig. 1D blue) (Moreno-Navarrete et al., 2013). Finally, loss of adipose triglyceride lipase or the mTOR pathway inhibitor TSC2 resulted in increased adipogenesis or SCREEN effects, which was entirely driven by a positive LIPID effect, consistent with increased lipid accumulation (Fig. 1D red) (Duvel et al., 2010; Schreiber et al., 2019). Thus, our screen not only identified candidate adipogenic regulators, but the MCE and LIPID scores provide testable hypotheses to understand how both known and novel adipogenic genes function during adipogenesis.

Of note, the physiological relevance of MCE to adipogenesis has been contended, in part because some cell line models of adipogenesis are capable of undergoing adipogenesis even in the presence of an inhibitor of mitosis. However, previous work has shown that these select cell line models will undergo endo-duplication in the presence of a mitotic inhibitor, and that inhibition of replication inhibits adipogenesis (Cho and Jefcoate, 2004). We confirm here that both replication and mitosis are required for adipogenesis of 3T3-L1 preadipocytes and primary human preadipocytes, supporting the value of studying both MCE and lipogenesis as key features of adipogenesis (Fig S1G).

### Assessment of screen performance

Having established the relationship between our three phenotypic scores, we next sought to assess our ability to detect known regulators of adipogenesis as well as MCE and LIPID. Focusing on specific gene targets first, inspection of our screen data showed that the known adipogenic regulators INSR, PIK3CA, C/EBPβ, PPARγ, KLF5, RB1, and ATGL scored as strong and significant regulators of adipogenesis (Fig. 2A, bottom). Further analysis showed that INSR, PIK3CA, C/EBPβ, SREBF2, and PPARγ score as significant inducers of MCE, while RB1 scores as a significant inhibitor of MCE (Fig. 2A top). Similarly, INSR, IRS1, PIK3CA, AKT2, RB1, and PPARγ score as significant inducers of LIPID, while ATGL scores as a significant inhibitor of LIPID (Fig. 2A middle). Thus, our screen detects well-established regulators of adipogenesis.

**Figure 2:**
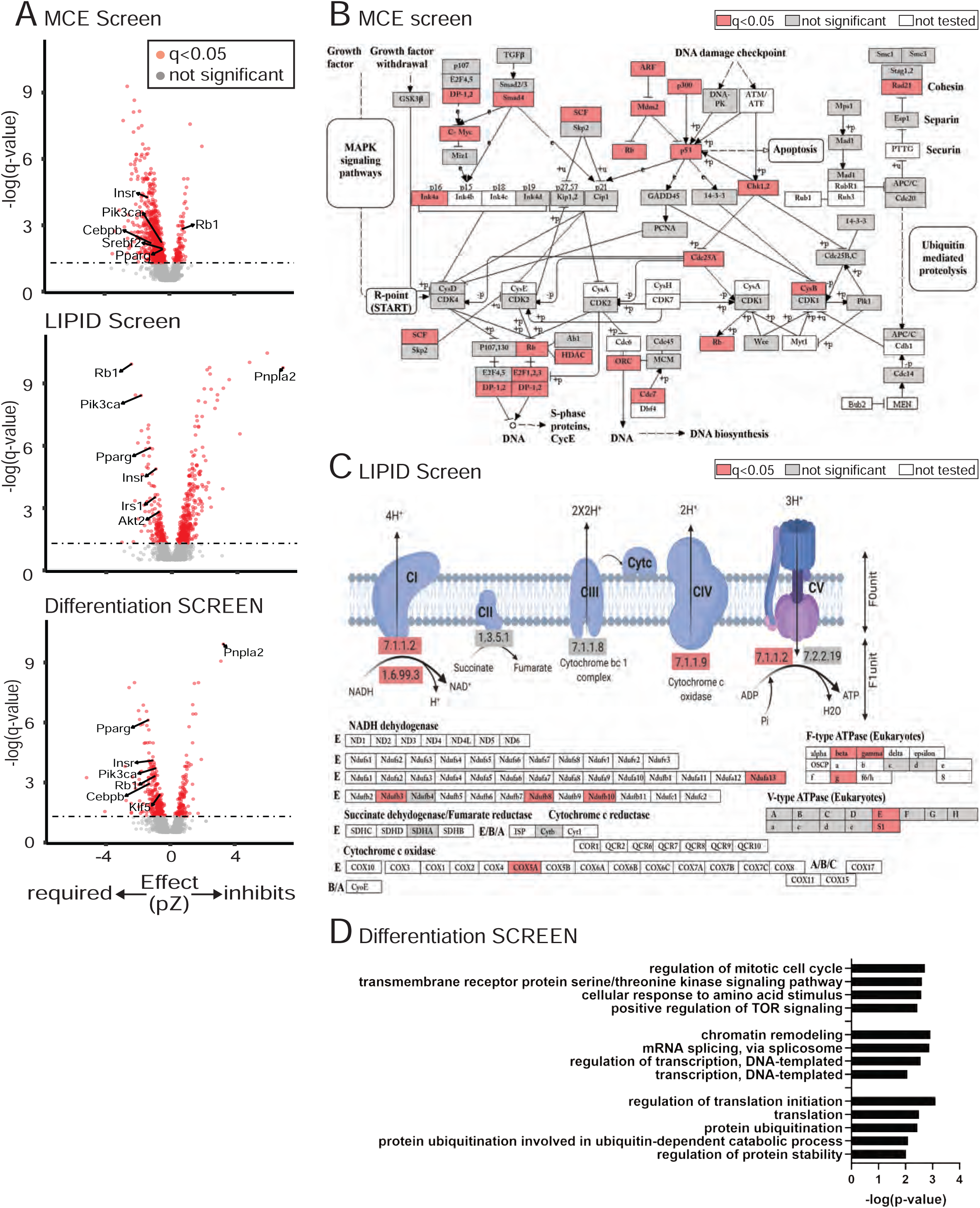
Overview of adipogenesis screen results. a. Volcano plot of q-value versus effect size for MCE, LIPID, and SCREEN. Known adipogenic regulators are labeled. b. KEGG pathway of cell cycle with gene targets that scored significantly in MCE analysis denoted in red. c. KEGG pathway of TCA cycle with gene targets that scored significantly in LIPID analysis denoted in red. d. GO term analysis of the SCREEN analysis identifies terms associated with signaling, regulation of transcription, and post-transcriptional regulation of protein translation and stability.

Next, we assessed the ability of our screen to recover hits in pathways known to regulate MCE and LIPID. Overlaying our MCE scores onto the KEGG pathway for the canonical cell cycle showed that many cell cycle genes were significant hits (Fig. 2B). This enrichment was observed even though many cell cycle genes were also required for viability and therefore dropped out of the screen. Similarly, overlaying our LIPID scores onto the KEGG pathways for the TCA cycle, FOXO signaling, and PI3K-AKT signaling, known to be important for lipogenesis and insulin signaling, showed that a considerable fraction of the components tested in these pathways were significant hits (Fig. 2C, Fig. S2A-B). Our screen thus identified components in diverse cellular processes, in accordance with the multiple pathways coordinating adipogenesis.

Finally, we performed a Gene Ontology Enrichment Analysis for biological processes on significant hits in SCREEN. Strikingly, all GO terms obtained fell into three general categories: signaling, regulation of transcription, and regulation of post-transcriptional processes (Fig. 2D). Adipogenesis is known to be regulated by a cascade of adipogenic transcription factors and co-regulators. Similarly, signal transduction pathways are known to regulate the initiation of adipogenesis and lipogenesis, including phosphorylation of key transcriptional regulators in response to initiation of adipogenesis (Humphrey et al., 2013; Rabiee et al., 2017). However, the prevalence of hits relating to the general regulation of translation and protein stability was striking, suggesting that these core machineries are integral to understanding adipogenesis.

### Many changes in protein abundance during adipogenesis are not driven by transcriptional mechanisms

Given the strong representation of hits in processes reflecting the central dogma for molecular biology, we next sought to assess the representation of our hits in the core machineries of transcription, post-transcriptional modifications, translation, and protein degradation (Fig. 3). Notably, this analysis not only underscored the importance of these gene families and complexes to MCE, LIPID, and SCREEN, but also highlighted the ability of this screen to identify which members of a gene family or molecular machinery are functionally most relevant. For example, many components of the translation initiation complex were significant hits, but specific isoforms of some components appeared exclusively. The scaffolding protein EIF4G consists of three family members. EIF4G3 is expressed at low levels and did not score in the screen. EIF4G1 is the major isoform and had dual, opposing roles in adipogenesis, being required for MCE yet inhibiting lipid accumulation. In contrast, the third isoform EIF4G2/DAP5 scored as the most significant inhibitor of adipogenesis and inhibited both MCE and lipid accumulation (Fig. S3). EIF4G2 is not well-understood, but generally thought to be responsible for approximately 20% of protein translation, notably including cap-independent translation (de la Parra et al., 2018). Thus, EIF4G1 and EIF4G2 have distinct functions during MCE, suggesting that the composition of the initiation complex may regulate translation of specific proteins with relevance to MCE. Similarly, analysis of cullin-ring complexes not only revealed that protein stability is critical in the regulation of adipogenesis, but also revealed that specific cullins, adaptor proteins, F-box, and other substrate recognition receptors are likely functionally important. This suggests that the degradation of specific target proteins is critical for adipogenesis. In addition, the specific targets of these degradation mechanisms are likely to have been identified as controllers of adipogenesis in our screen.

**Figure 3:**
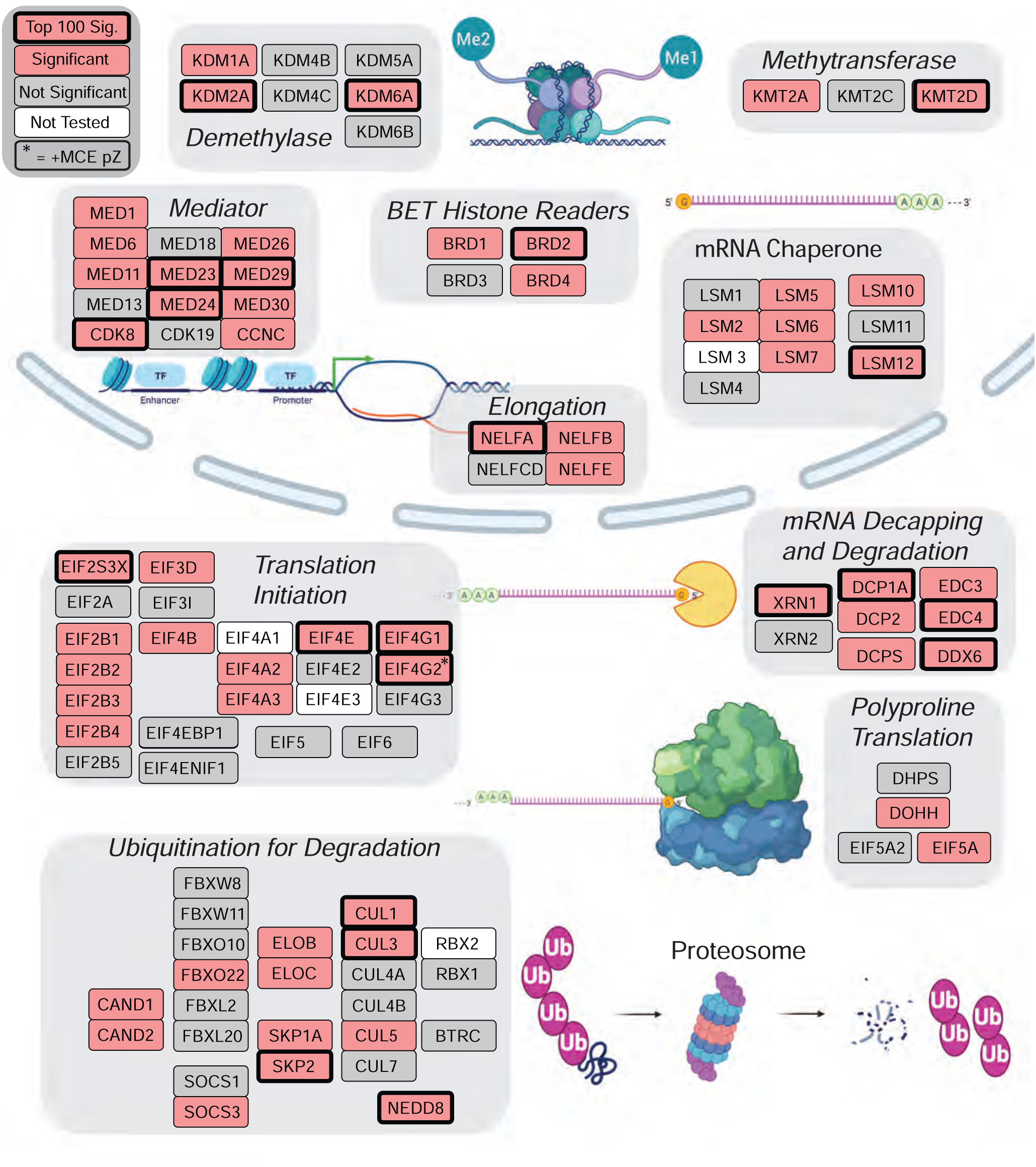
Complexes Important to Adipogenesis. Gene families and complexes important for core processes in the central dogma for molecular biology are strongly represented among hits. Light red color indicates the genes that had a significant score (q<0.05) for MCE, LIPID, or SCREEN. Border denotes a significance score in the top 100 in MCE, LIPID, or SCREEN and * denotes a positive pZ in MCE. This analysis highlights which subunit or gene family member is critical for adipogenesis.

There have been numerous gene expression profiles conducted to assess changes in gene expression during 3T3-L1 adipogenesis to identify candidate adipogenic regulators (Ahmed and Kim, 2019). However, our data argued that processes regulating protein abundance directly, namely differential translation of mRNA and degradation of proteins, are also critical for adipogenesis. Therefore, to gain a clearer picture of the diverse mechanisms that drive adipogenesis, it is critical to move beyond the well-established role of transcription in adipogenesis and consider whether specific translational control mechanisms may contribute. We might imagine specific proteins, identified in our screen as important for adipogenesis, that showed strong changes in protein without notable change in mRNA or where both protein and mRNA show strong changes.

We first took a proteogenomic approach and sought to correlate changes in mRNA and protein concentrations during 3T3-L1 adipogenesis. Specifically, duplicate samples at 0, 24, 48, and 96 hours post-differentiation were collected and analyzed by shotgun proteomics to quantify changes in protein abundance during 3T3-L1 adipogenesis. (Fig. 4A, Table S3). Newly obtained and previously published gene expression data for the same time points were combined to obtain a consensus of mRNA expression during 3T3-L1 adipogenesis (Fig. S4A). Scatter plot analysis showed that there is very poor overall correlation between mRNA and protein abundance at any of the time points assessed. In fact, this analysis shows that only approximately 25% of protein abundance can be explained by mRNA concentration (Fig. 4B). Numerous studies have shown that the relationship between protein and mRNA is complicated by the variable contribution of other processes, including regulatable translation rates, protein stability, and protein secretion (Liu et al., 2016). However, the correlation observed in this study is even poor compared to the more typically observed squared Pearson correlation coefficient of ∼0.40 (Vogel and Marcotte, 2012). To determine how changes in mRNA expression related to changes in protein abundance over the time course of adipogenesis, we next performed hierarchical clustering of the proteomics time course data (Fig. 4C, Table S4). This analysis yielded 10 clusters with differential changes in protein expression during adipogenesis. Next, we assessed mRNA expression changes for the proteins found in each cluster by performing hierarchical clustering (Fig. 4D, Fig. S4B, Table S4). A number of specific patterns linking gene expression to protein abundance emerged from this analysis. Some proteins accumulate concurrently with mRNA expression (e.g. Fig. 4D cluster 2.A). Other proteins accumulate appreciably later than mRNA, suggesting that either translational control or regulation of protein stability could control accumulation of proteins important for differentiation (e.g. Fig. 4D cluster 7.A). Finally, many of the transcript changes bore little to no resemblance to the changes in protein concentration (e.g. Fig. 4D cluster 3.B). For example, both TSC2 and NCOA6 protein abundance transiently increase during adipogenesis, while mRNA expression remains unchanged or transiently decreases during adipogenesis (Fig. 4D cluster 3.B, Table S4). This suggests that proteins are selectively translated (or selectively stabilized) upon addition of differentiation cocktail and then specifically degraded upon completion of MCE. Consistent with its known function as an mTOR pathway inhibitor, TSC2 inhibited LIPID in our screen (Table S2). The expression pattern suggests that TSC2 protein is transiently upregulated during early adipogenesis to prevent premature accumulation of lipids. Similarly, NCOA6 is a known co-activator of LXRs and critical for liver lipid metabolism (Kim et al., 2003). Moreover, it was identified in a GWAS study for body fat distribution (Heid et al., 2010; Winkler et al., 2015). In our screen, NCOA6 is required for LIPID (Table S2). The expression pattern suggests that NCOA6 protein is upregulated during early MCE to initiate transcriptional expression of genes important for lipogenesis. Finally, NUBP2 protein transiently decreases at the 48h time point, when mRNA levels are maximal (Fig. 4D cluster 4.B, Table S4). We found that NUBP2 is a novel regulator of adipogenesis, required for MCE (Fig. S1B, Table S2). Since NUBP2 is required for the formation of iron-sulfur clusters and hence proliferation (Christodoulou et al., 2006), the expression pattern during adipogenesis suggests that NUBP2 may be degraded at the end of MCE to enable cell cycle exit. Additional proteins that are strongly controlled at the level of protein abundance and important for adipogenesis were found to be highly linked to GWAS studies supporting roles in obesity (such as body mass index), including MED11, TAOK1, FNT, and WDR82 (Table S2, Table S4). Other proteins were linked to metabolic disorders. Thus, we conclude that focusing on changes in protein abundance, independent of mRNA expression, may identify interesting adipogenic regulators. An examination of conserved processes regulating protein abundance may uncover key mechanisms regulating adipogenesis. Here, we focus on the importance of hypusination, a highly conserved pathway regulating translation of proteins containing consecutive proline residues, and neddylation, a ubiquitin-like posttranslational modification known to activate Cullin-Ring E3 ubiquitin ligases.

**Figure 4:**
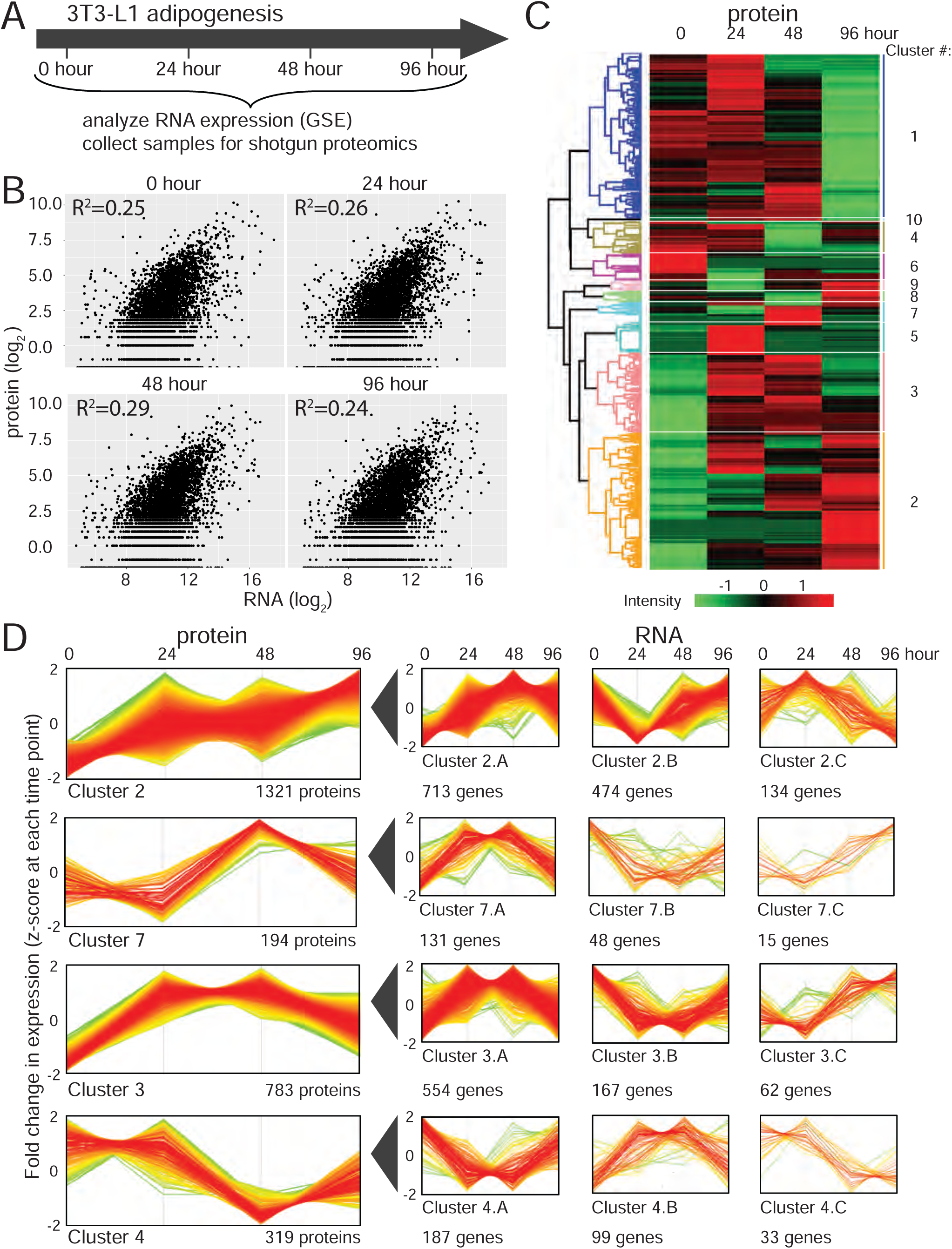
Protein translation and stability drive substantial dynamic changes in expression during adipogenesis. a. Schematic for analysis of RNA and protein abundance during 3T3-L1 adipogenesis. RNA abundance was obtained from published data, protein abundance was determined using shot-gun proteomics. b. Scatter plot showing log_2_ of RNA versus protein abundance at 0, 24, 48, and 96 hours of 3T3-L1 adipogenesis. Only 25% of protein abundance can be explained by mRNA abundance. c. Heatmap showing hierarchical clustering of protein abundance during 3T3-L1 adipogenesis. Changes in protein concentration fall into 10 clusters. d. Profile plot showing fold change in protein expression over time (after z-scoring for each time point) for four clusters. For all proteins in each cluster, corresponding RNA concentrations were analyzed, and hierarchical clusters yielded three profile plots. Number of proteins/genes in each cluster is indicated below each cluster. Color of profile denotes distance from center, where red is zero and green is maximal change.

### Hypusination is required to translate proline-rich and other adipogenic regulators important for MCE

Examination of the screen showed that proteins important for the translation initiation are important for adipogenesis (Fig. 3). Additionally, the screen identified that the highly conserved pathway for protein hypusination, generally linked selectively to translation initiation also scored highly. Hypusine is a unique post-translational modification of the EIF5A translation initiation factor, and it is essential for EIF5A function (Fig.5B). Importantly, EIF5A is important for the efficient translation of specific mRNAs, including those encoding four or more consecutive proline codons (Gutierrez et al., 2013), however eIF5A may sustain other forms of specialized translation. There are two isoforms, the ubiquitously expressed isoform EIF5A and the testes and brain-specific isoform EIF5A2 (Jenkins et al., 2001). In a highly conserved pathway, the enzymes deoxyhypusine synthase (DHPS) and deoxyhypusine hydroxylase (DOHH) use the polyamine spermidine as a substrate for hypusination of a conserved lysine residue in EIF5A (Abbruzzese et al., 1986; Park et al., 1981; Wolff et al., 1995).

**Figure 5:**
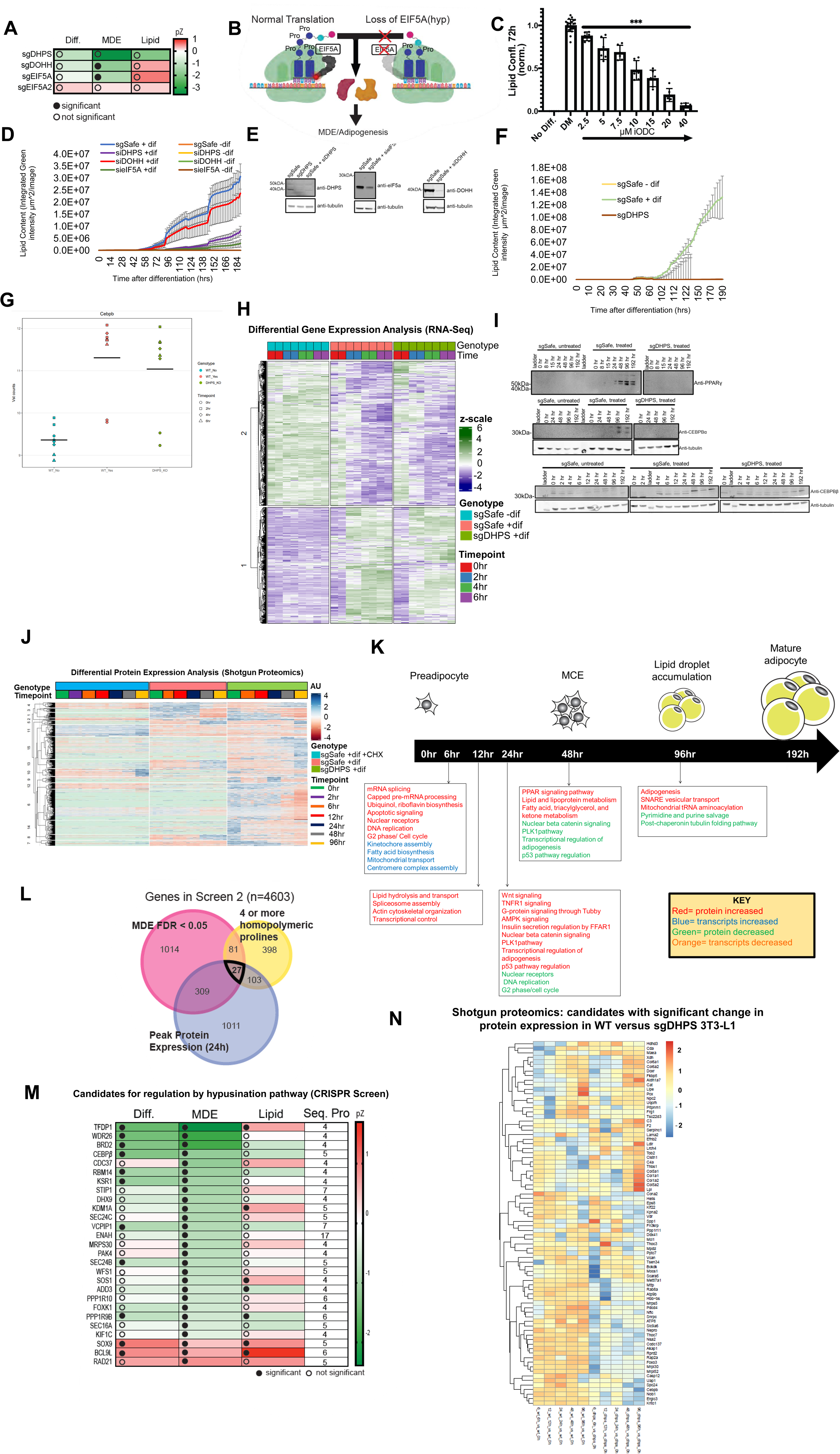
Hypusination Pathway is critical for mitotic clonal expansion and adipogenesis. a. Heatmap displays phenotypic scores of components of the hypusination pathway. Loss of DHPS, DOHH, and EIF5A inhibits MCE, EIF5A2 does not. b. A model for the hypusine-eIF5A high efficiency translation initiation pathway. The hypusine-eIF5A is thought to be important for high efficiency translation, especially important for difficult to synthesize protein sequences contraining tracts of multiple prolines. c. Treatment of 3T3-L1 preadipocytes with iODC (DFMO), a hypusination pathway inhibitor, inhibits adipogenesis in a dose-dependent manner as measured by BODIPY staining of lipid content. Mean of two experiments +/- SD. ***p < 0.001. d. siRNA knockdown of DHPS, DOHH, and eIF5A translation initiation components shows deficiency in adipogenesis by BODIPY staining of lipid content. e. Validation of DHPS knockout by immunoblot analysis. f. CRISPR-Cas9 knockout of DHPS translation initiation regulator shows deficiency in adipogenesis by BODIPY staining of lipid content. g. CEBP/b transcription is induced by differentiation cocktail, but unaffected by DHPS knockouts h. RNASeq expression profiling of 3T3-L1 cells with or without differentiation cocktail and DHPS knockout shows the hypusination-translation initiation pathway has minimal effect on the induction of mRNAs during adipogenesis. i. DHPS knockouts show substantial reduction in accumulation of key transcription factor markers of differentiation j. Shotgun proteomic analysis of the kinetics of protein expression during 3T3-L1 adipogenesis demonstrates selective translation of critical factors. MORE. k. GO analysis of critical classes of factors translated at specific times during adipogenic differentiation l. Venn Diagram to identify candidate regulators of MCE whose translation is dependent on the hypusination pathway. Candidates are regulators of MCE, contain a stretch of four or more prolines and are expressed during MCE. This analysis identifies 27 candidate regulators m. Heatmap displays phenotypic scores of candidate regulators and the longest stretch of prolines in sequence. This list contains a number of proteins known to be important for adipogenesis and specifically MCE. n. Clustering translationally induced factors lost under DHPS knockout supports a broad program of differentiation factors under translational control

In our initial validation of the first screen, we confirmed that loss of DHPS inhibited adipogenesis (Fig. S1B). The second screen identified DOHH and EIF5A, but not EIF5A2, as required for MCE (Fig. 5A, Fig. S5A). Of note, loss of DHPS also resulted in a very negative MCE score, but this did not reach the significance cut-off, likely because more than half of the targeting sgRNAs were not recovered in any of the experimental time points, consistent with loss of DHPS affecting cell viability (Lee et al., 2002). We used siRNA to knock down DOHH, DHPS, and EIF5A in 3T3-L1 to test the hypothesis that the hypusination pathway promotes adipogenesis (Figure 5D), confirming the efficacy of knockdown via Western (Figure 5E). After knockdown, cells were treated with standard differentiation cocktail (IBMX + insulin + dexamethasone) to induce adipogenesis and stained with BODIPY lipid droplet live cell imaging dye. Cells were imaged every two hours over the course of 8 days to track adipogenesis over time using a the Incucyte imaging system. All three knockdowns demonstrated a strong inhibition of adipogenesis compared to WT 3T3-L1 after treatment with differentiation cocktail, though with varying degrees of severity. Interestingly, knockdown of DHPS showed the strongest inhibition of adipogenesis, followed by EIF5A and DOHH (Figure 5D), perhaps suggesting the influence of unknown factors in at least partially rescuing the adipogenesis defects in EIF5A and DOHH knockdown cells. As the strongest candidate out of the three genes, we further validated the role of DHPS by using CRISPR-Cas9 to generate stable cell lines with frameshifting mutations early in the coding sequence of DHPS. The goal was to generate cell lines encoding early stop codons in DHPS that would therefore yield non-functional protein products (See Supplement for guides). We validated the loss of protein DHPS via Western blot (Fig. 5E). Because of the possibility that there may still be protein product which could technically make the cell line not a true “knock out” (though likely not functional), we will refer to these cells as “hypomorphic mutants.” We also generated sgEIF5A and sgDOHH 3T3-L1 cell lines which also showed the same trends as the siRNA data (not shown). The DHPS hypomorphic mutants mirrored knockdown data and also showed a strong inhibition of adipogenesis (Fig. 5F).

To further validate the importance of hypusination to MCE, we tested if acute pharmacological inhibition of this pathway reduced 3T3-L1 adipogenesis. Specifically, we treated 3T3-L1 cells with an inhibitor of ornithine decarboxylase (ODC1), the rate-limiting enzyme in spermidine synthesis (Meyskens and Gerner, 1999). The inhibitor was added two days prior to addition of differentiation cocktail to sufficiently deplete the spermidine pool prior to MCE and the inhibitor was maintained with the cocktail. Inhibition of hypusination resulted in a dose-dependent inhibition of MCE and adipogenesis (Fig. 5C, Fig. S5B). Of note, spermidine has previously been shown to be required for 3T3-L1 adipogenesis (Vuohelainen et al., 2010). Our data argue that loss of spermidine inhibits adipogenesis at least in part by inhibiting hypusination.

To verify that the hypusination exclusively drives translational control versus transcriptional control, we performed RNA-seq and shotgun proteomics of WT versus sgDHPS 3T3-L1. In comparing the transcriptional profile of sgDHPS versus WT preadipocytes with and without induction with differentiation cocktail in the first 12hrs, we observe that sgDHPS cells had a transcriptional profile that more closely mirrored differentiated versus undifferentiated 3T3-L1 (Fig.5 G-I). In looking at the transcript levels of known regulators of adipogenesis such as C/EBPβ, we note that there is no significant difference between WT differentiated versus sgDHPS. These data support the model that the hypusination pathway regulates adipogenesis independent of transcriptional control. Shotgun proteomics, on the other hand, revealed a striking difference in protein levels between WT and sgDHPS cells treated with differentiation cocktail. We verified that loss of sgDHPS does not lead to universal loss of translational control, as cycloheximide-treated 3T3-L1 demonstrated near universal inhibition of translation whereas sgDHPS showed inhibition only for a select pool of translated proteins (Fig. 5K). Altogether, the mapping of transcripts and protein products during adipogenesis revealed to us striking new patterns of the diverse signaling events coordinating preadipocyte fate change (Fig. 5L).

In one model, we hypothesized that hypusination is required for MCE because hypusinated EIF5A is required for the translation of a subset of mRNAs that code for long sequences of consecutive prolines and that one or more of these gene products is in turn required for MCE (Fig. 5B). We therefore sought to generate a list of candidate adipogenic proteins containing multiple consecutive prolines (Fig. 5M). Specifically, we sought genes that were identified in the genome-wide CRISPR screen as regulating MCE (Fig. 5M red circle), containing four or more consecutive prolines (yellow circle), and which are translated during MCE (blue circle). This analysis identified 27 candidates (Fig. 5N). Notably, this list contains both well established and candidate adipogenic genes. C/EBPβ is a known early adipogenic transcription factor required for MCE (Tang et al., 2003a). Consistent with our model, a previous study has shown that loss of spermidine prevents the translation, but not the transcription, of C/EBPβ in early adipogenesis (Hyvonen et al., 2013). Other candidate proteins are also required for adipogenesis, including RBM14 (Firmin et al., 2017) and BRD2 (Denis et al., 2010; Wang et al., 2009), but their importance to MCE is less understood. Thus, we proposed that the hypusination pathway regulates adipogenesis by enabling the translation of regulators of MCE with multiple consecutive prolines, including C/EBPβ. In support of this model, we note that sgDHPS CRISPR cell lines showed depleted C/EBPα, C/EBPβ, and PPARy protein expression via Western blot (Fig.5J). We then turned to our shotgun proteomics data to see if loss of DHPS led to a significant decrease in translation of proteins with long proline stretches. Upon closer examination, though, we were interested to see that of the 143 proteins that were upregulated in WT but not sgDHPS 3T3-L1. The 143 proteins regulate a number of critical functions in adipogenesis, based on positive hits from our screen or known regulators. These include cell division, endocytosis and vesicular transport, transcription, and mitochondrial and cytosolic ribosomal assembly (Fig. 5O). We believe that many of these are proteins that must be selectively translated to achieve efficient adipogenesis. Whether the hypusination pathway drives these critical regulators of early adipogenesis by enhancing selective translation of long stretches of prolines or by other mechanisms remains to be elucidated. A recent study has highlighted the genetic importance of translation initiation in *de novo* adipogenesis [ref. Ruggero paper PMID 33619378]. Based on the broad effect of translation initiation factors and the hypusine regulatory pathway, coupled with our data establishing selective translation of essential factors for adipogenesis, we propose that these 143 factors provide a first draft of the selective translation program for adipogenesis.

### Neddylation is required to prevent precipitous lipid accumulation in preadipocytes and hepatocytes

Examination of our screen showed that the ubiquitin-like protein NEDD8 was a highly significant hit (Fig. 6A, top row). We were particularly intrigued by the dual role of NEDD8 in adipogenesis. While NEDD8 was required for adipogenesis, this was driven by a requirement for MCE, consistent with previous reports (Dubiel et al., 2017; Park et al., 2016). Our screen argued that NEDD8 also independently inhibits lipid accumulation. NEDD8 is a ubiquitin-like post-transcriptional modification and its best validated and central substrate is the cullin subunit of the Cullin-Ring E3 family of ubiquitin ligases (CRLs) (Petroski and Deshaies, 2005). Neddylation is required to activate CRLs, which in turn promotes ubiquitin-mediated degradation of their substrates. CRLs are multi-subunit complexes containing a cullin, a RING-box protein, an adapter, and a specific substrate recognition receptor. There are eight cullins, two RING-box proteins, 4 adapter proteins, and a multitude of substrate recognition receptors (including F-box proteins), making this the largest superfamily of E3 ubiquitin ligases. Consequently, CRLs are responsible for ubiquitination of ∼20% of proteins targeted for ubiquitin-mediated degradation and play a critical role in many cellular processes (Zhao and Sun, 2013). The best-established function for CRLs is the regulation of the cell cycle and consistent with this, we observed a significant negative MCE score in the screen (Fig. 6A, top row). Of the eight cullin genes, six were tested in the second screen. Strikingly, examination of their scores allowed for the identification of which cullins and other CRL components functioned in MCE versus LIPID. Based on the screen, neddylation of CUL3 is required for MCE, consistent with published data (Fig. 6A, second row) (Dubiel et al., 2017). In contrast, the screen shows that neddylation of CUL1 or CUL5, as well as their association with the CUL5 adapter proteins ELOB and ELOC, is required for inhibiting lipid accumulation (Fig. 6A). We decided to further explore how neddylation of this CRL regulates lipid accumulation.

**Figure 6:**
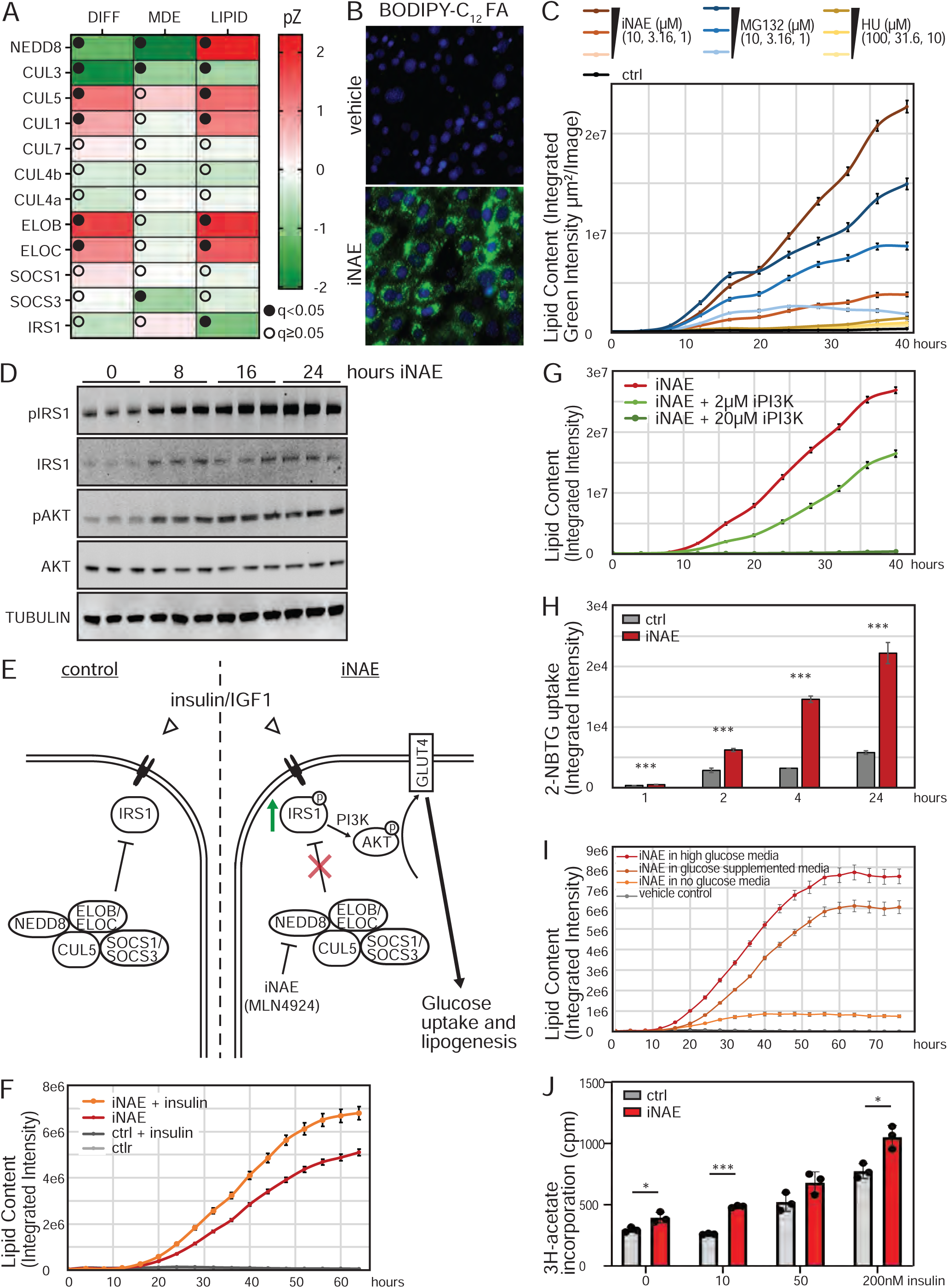
Neddylation Pathway inhibits premature accumulation of lipids. a. Heatmap displays phenotypic scores of NEDD8 and Cullin-RING Ubiquitin ligase complexes. Neddylation of some CRLs is required for adipogenesis, likely driven by the requirement for MCE. Loss of neddylation also promotes lipid accumulation via a distinct CRL. b. 3T3-L1 preadipocytes treated for 32 hours with an inhibitor of neddylation accumulate the green fluorescent BODIPY-C_12_ FA. c. 3T3-L1 preadipocytes treated with an inhibitor of neddylation or with an inhibitor of proteasome degradation, but not with an inhibitor of proliferation, accumulate lipid in a dose-dependent manner as measured by BODIPY fluorescent intensity. Data points represent average of 3 independent experiments ±SE. d. Inhibition of neddylation in 3T3-L1 preadipocytes promotes activation of the insulin signaling pathway as determined by immunoblotting for phosphorylated and total IRS1 and AKT protein. Immunoblot shows three independent experiments per time point e. Model describing how neddylation inhibits *de novo* lipogenesis. In the absence of neddylation, components of the insulin signaling pathway are stabilized, leading to activation of the pathway and lipid accumulation f. Insulin potentiates accumulation of lipids induced by inhibition of neddylation in 3T3-L1 preadipocytes as measured by BODIPY fluorescent intensity. Data points represent average of 3 independent experiments ±SE. g. Inhibition of PI3K attenuates the accumulation of lipids promoted by inhibition of neddylation in a dose-dependent manner as measured by BODIPY fluorescent intensity. Data points represent average of 3 independent experiments ±SE. h. Inhibition of neddylation in 3T3-L1 preadipocytes promotes accumulation of the fluorescent glucose analog 2-NBTG. Bar graphs are average of 3 independent experiments ±SD. *** p<0.001; i. Inhibition of neddylation in 3T3-L1 preadipocytes only promotes lipid accumulation in the presence of glucose, consistent with *de novo* insulin-induced lipogenesis. Data points represent average of 3 independent experiments ±SE. j. Inhibition of neddylation promotes lipogenesis in AML12 hepatocytes as determined by incorporation of [3H]-acetate. Insulin potentiates this effect. Bar graphs are average of 3 independent experiments ±SD. * p<0.05; *** p<0.001;

First, we sought to confirm that inhibition of neddylation resulted in increased lipid accumulation. Specifically, 3T3-L1 preadipocytes were treated with a specific inhibitor of NAE, the E1 ubiquitin activating enzyme in the neddylation pathway (iNAE, MLN4924). No differentiation cocktail was added. Lipid content was assessed using the lipophilic dyes BODIPY or LipidTOX. Within 2 days, inhibition of neddylation led to the formation of many small lipid droplets in 3T3-L1 preadipocytes (Fig. S6A, top panels). The time frame of this lipid accumulation as well as the disorganized appearance of the lipid droplets argued against this being a consequence of canonical adipogenesis. Consistently, iNAE promoted lipid accumulation even in the presence of an inhibitor of PPARγ (Fig. S6B). Co-staining with BODIPY and PLIN, a protein upregulated during adipogenesis to coat lipid droplets, confirmed that the intracellular lipid accumulated in response to neddylation inhibition was not organized into the typical lipid droplet known to form in differentiating preadipocytes (Fig. S6A, bottom panels). To further study this intracellular lipid, 3T3-L1 preadipocytes were grown in the presence of a fluorescently labeled fatty acid (FA) with and without iNAE (Fig. 6B). In the absence of neddylation, 3T3-L1 preadipocytes accumulate lipids precociously.

Since neddylation of CRLs is required for ubiquitin-mediated degradation of substrates, including cell cycle proteins, we next sought to determine if lipid accumulation was a consequence of cell cycle arrest or the stabilization of other CRL substrates. 3T3-L1 preadipocytes were treated with inhibitors of neddylation, the proteasome (MG132), or S-phase (hydroxyurea, HU). As expected, inhibition of neddylation or S-phase inhibited proliferation in a dose-dependent manner (Fig. S6C). Only inhibition of neddylation or the proteasome, and not cell cycle inhibition, resulted in lipid accumulation in a dose-dependent manner, as assessed by green-fluorescent BODIPY staining over time (Fig. 6C). This lipid accumulation was observed within hours of inhibition. Since inhibition of the proteasome phenocopied the effect of loss of neddylation, we hypothesized that a neddylated (and hence active) CRL constitutively promotes proteasomal degradation of a substrate which normally promotes lipid accumulation. A candidate for this substrate would be stabilized in the absence of neddylation. To identify proteins stabilized in the absence of neddylation, 3T3-L1 preadipocytes treated with iNAE or vehicle were analyzed by shotgun-proteomics (Table S5). PANTHER pathway analysis of differential protein abundance revealed a significant enrichment for cell cycle regulators, consistent with the known role of CRLs in regulating the cell cycle (Fig. S6D, orange and teal). Notably, proteins involved in insulin signaling were also among the top hits, including the insulin signaling adapter IRS1 (Fig. S6D, purple and teal). IRS1 has previously been identified as a substrate for CRLs in other cell contexts (Hu et al., 2012; Scheufele et al., 2014; Xu et al., 2012). Immunoblot staining confirmed that inhibition of neddylation resulted in activation of insulin signaling within hours of treatment, as assessed by staining for phosphorylated IRS1 and AKT (Fig. 6D). Consistent with the proteomics data, this is at least in part due to increased stabilization of IRS1 protein upon neddylation inhibition. In contrast, total AKT protein levels are unchanged, arguing that AKT is not a CRL substrate and that activation of the pathway is instead driven by stabilization of other insulin signaling pathway components, notably IRS1. Thus, we propose that in 3T3-L1 preadipocytes the neddylated CRL consisting of CUL5, ELOB/ELOC and SOCS1 and SOCS3 targets components of the insulin signaling pathway, including IRS1, for degradation. Consistent with this, our screen confirmed that IRS1 is required for lipid accumulation (Fig. 6A, last row). In the absence of neddylation, components of the insulin signaling pathway, including IRS1, would thus accumulate, resulting in uncontrolled insulin signaling and lipogenesis (Fig. 6E).

To test this model, 3T3-L1 preadipocytes treated with an inhibitor of neddylation were supplemented with additional insulin. This potentiated lipid accumulation in the presence of iNAE (Fig. 6F). Similarly, addition of an inhibitor of PI3K signaling, required for AKT activation, prevented iNAE-induced lipid accumulation in a dose-dependent manner (Fig. 6G). Next, we tested if inhibition of neddylation resulted in increased *de novo* lipogenesis. First, we assessed the ability of 3T3-L1 preadipocytes treated with the inhibitor of neddylation to take up glucose, the macromolecule commonly catabolized to generate the building blocks for FA synthesis. Addition of iNAE resulted in a dramatic and highly significant increase in the uptake of the fluorescent glucose analog 2-NBTG (Fig. 6H). Moreover, inhibition of neddylation only resulted in lipid accumulation if 3T3-L1 cells when cultured in the presence of glucose (Fig. 6I). Finally, we tested whether loss of components of the CRL would mimic and/or synergize with iNAE-induced lipid accumulation. Knockdown of EloB or double-knockdown of Socs1 and Socs3 promotes lipid accumulation in 3T3-L1 cells, enhanced in the presence of the neddylation inhibitor (Fig. S6E). Moreover, loss of IRS1 inhibited iNAE-induced lipid accumulation, supporting our model that iNAE-induced stabilization of IRS1 is a key driver of lipid accumulation (Fig. S6E). Together, our model proposes that iNAE-induced lipid accumulation is the consequence of insulin signaling-induced glucose catabolism to synthesize intracellular lipids *de novo*.

Finally, we wanted to assess if this role of neddylation is conserved in other cell contexts sensitive to insulin-induced lipogenesis. Notably, preadipocytes isolated from gonadal white adipose tissue, but not from inguinal white adipose tissue, showed iNAE-induced lipogenesis in culture both in the presence and absence of differentiation cocktail (Fig. S6F and data not shown). Since gonadal preadipocytes show little adipogenesis in culture even in the presence of differentiation cocktail while inguinal preadipocytes differentiate spontaneously even in the absence of differentiation cocktail, this further supports the hypothesis that neddylation is required to specifically prevent precocious activation of insulin signaling and lipogenesis. Similarly, inhibition of neddylation promotes lipid accumulation in another cell line model of adipogenesis, C3H10T1/2 cells, in a dose-dependent manner (Fig. S6G). Finally, we investigated the importance of neddylation in hepatocytes. Non-alcoholic fatty liver disease is strongly correlated with obesity and in part driven by increased hepatic lipogenesis (Divella et al., 2019). Inhibition of neddylation in AML12 hepatocytes resulted in a significant increase in *de novo* lipogenesis, further potentiated by insulin in a dose-dependent manner (Fig. 6J). Thus, neddylation is required to prevent precocious, insulin-induced lipogenesis in diverse cell populations.

## Discussion

Obesity-related diseases are among the leading causes of morbidity and preventable death. Identifying regulators of adipogenesis may advance several potential therapeutic avenues. Here, we present a genome-wide screen for regulators of adipogenesis, both validating well-characterized regulators of adipogenesis and identifying novel candidates. We also quantify the relative importance of adipogenic hits by MCE and lipogenesis. To illustrate the power of the screen, the screen uncovers that hypusination is specifically required for translating inducers of MCE and that neddylation is required to prevent premature, ectopic lipogenesis in both preadipocytes and hepatocytes. This follow-up analysis highlights the importance of post-transcriptional mechanisms to the regulation of adipogenesis and expands the focus on transcriptional control as the pivotal regulator of adipogenesis.

### Resources provided in this study

The functional genomics screen enumerates many candidate adipogenic regulators. To further aid future investigators, we list as an additional parameter which hits were identified in a GWAS study for body fat distribution (Table S2) (Heid et al., 2010; Winkler et al., 2015). Moreover, this study also provides a compilation of public gene expression data during 3T3-L1 adipogenesis. Finally, we performed a proteomic analysis to quantify changes in protein abundance during 3T3-L1 adipogenesis. We used publicly available mRNA expression data and performed a proteogenomic comparison of 3T3-L1 adipogenesis (Table S4).

A main conclusion uncovered in our analysis is that mechanisms regulating translation of subsets of mRNA and protein stability play a critical role in adipogenesis. Notably, the screen provides testable hypotheses for how these post-transcriptional mechanisms function during MCE and lipogenesis, as illustrated for hypusination and neddylation. For example, our data showed that a translation initiation complex containing EIF4G1 is required for MCE, while translation initiation utilizing EIF4G2/DAP5 inhibits MCE and adipogenesis. Previous studies have shown that EIF4G2 drives translation of a select group of mRNAs important for diverse cellular processes, including the transition of mESCs from pluripotency to differentiation (Sugiyama et al., 2017; Yoffe et al., 2016). Additional teratogenesis studies with mESCs lacking EIF4G2 have demonstrated a requirement for all tissues of mesodermal origin, further implicating this gene in the earliest stages of embryonic differentiation (Yamanaka et al., 2000). These studies suggest that EIF4G1 and EIF4G2 selectively translate proteins that are either required for or are inhibitory to MCE, respectively and that these roles are vital for proper mesoderm differentiation. Further investigation may discover the identity of these adipogenesis-associated mRNAs.

Finally, while much of the screen validation in this study focused on post-transcriptional control of adipogenesis, we note that this screen also identified a number of candidate adipogenic transcription factors, including Zmynd8, which has been linked by GWAS studies to body fat distribution (Table S2). Other significant hits were associated with the cytoskeleton and mechanisms of cellular trafficking. Moreover, the screen also identified candidate adipogenic genes that are annotated but are uncharacterized to-date, including GM14393 and E130309D02Rik (Fig. S1C). Follow-up investigation will establish how these genes regulate steps in adipogenesis and lipogenesis and may suggest potential therapeutic targets for obesity and metabolic health.

### Limitations of the screen

To avoid the confounding effects of viability genes in this screen, the infected pool of 3T3-L1 cells was passaged for two weeks prior to initiating the experiment (defined as time of plating, 4 days prior to Day 0). This time frame was selected to (1) allow for sufficient turnover of target genes to disrupt gene function and (2) select against any sgRNA targeting viability genes, since a decrease in sgRNA abundance over time due to loss of viability could be interpreted as a requirement for adipogenesis and MCE. While the two-week time frame should be sufficient to select against sgRNA with strong lethality, we cannot exclude the presence of some sgRNA in the pool that resulted in a slight decrease in cell growth over time. However, since the average recovery of sgRNA elements per gene for the duplicates was 19.34 out of 20 sgRNA total, we do not believe that this is a major concern. Likewise, some genes required for viability may have additional, but separable roles, in adipogenesis which we would fail to identify here.

The screen is based on the ability to enrich for cells with high or low amounts of intracellular lipid by sorting for green fluorescent intensity upon addition of the lipophilic dye BODIPY. However, if a cell is infected with a sgRNA that substantially affects total cell number, then this cell may be enriched in the BODIPY^high^ or BODIPY^low^ cell populations independent of adipogenesis. We observed this effect for components of the Hippo signaling pathway. Absence of Hippo signaling results in an inability to arrest at confluency and to exit MCE. Since the 3T3-L1 adipogenesis protocol specifies that cells remain confluency-arrested for two days prior to addition of differentiation cocktail and that cells normally exit MCE after 2 days of differentiation, loss of Hippo signaling effectively resulted in four additional days of proliferation compared to cells infected with non-targeting sgRNAs. This yielded a positive SCREEN score due to increased representation of sgRNAs in the Day 4 cell population (Table S1), despite the fact that Hippo signaling is required for adipogenesis and consequently leads to decreased lipid content (data not shown) (Ardestani et al., 2018). We cannot exclude the possibility that other pathways regulating confluency are similarly over/underrepresented, confounding the analysis. Therefore, data related to these pathways should be viewed with caution. We note, however, that the screen was sufficiently robust to screen cell cycle regulators, including the RB1/E2F signaling pathway. Similarly, if a cell is infected with a sgRNA that affects lipid content too dramatically, then a true LIPID score may drive a significant SCREEN score, without in fact affecting adipogenesis. We observed this effect for PNPLA2/ATGL, which had the single highest score in LIPID. ATGL is known to regulate lipolysis, consistent with its LIPID score. However, mouse ATGL KO mice have normal adipogenesis in their fat cell populations, despite having a significant SCREEN score here. We note that the scores, and the relationship between the scores, hold true for most cases and only falls short for extreme phenotypic scores, such as the LIPID score for PNPLA2/ATGL.

Finally, we have taken advantage of the fact that differentiated adipocytes accumulate lipid by using lipid content as a marker for differentiation. Follow-up validation of hits identified in the screen should test additional markers of adipogenesis in real-time (e.g. activation of PPARγ). Despite the limitations of our CRISPR genome-wide screen, this data set will lay the foundation for identifying and analyzing understudied mechanisms driving adipogenesis, paving the way for many future studies.

### Homopolymeric proline proteins driving MCE

Our screen showed that the highly conserved hypusination pathway, required for translating proteins with sequences of consecutive prolines, is important for MCE. We generated a list of candidate drivers of MCE that contain a sufficiently long sequence of prolines to require hypusinated EIF5A for efficient translation. Of note, the murine C/EBPβ sequence contains five consecutive prolines, and spermidine depletion is known to inhibit translation of C/EBPβ while not affecting mRNA expression (Hyvonen et al., 2013). Translation of C/EBPγ, which functions similarly to C/EBPβ but does not contain this stretch of prolines, is not affected by spermidine depletion (Hyvonen et al., 2013). Murine C/EBPα contains a stretch of eight prolines, and therefore also likely requires the hypusination pathway for efficient translation, albeit later in the adipogenic program. Importantly, human C/EBPβ and C/EBPα also contain homopolymeric proline sequences with nine and seven consecutive prolines, respectively. Thus, the hypusination pathway is likely also critical for adipogenesis of human preadipocytes.

We further validated that hypusination is a critical driver of translation not transcription using a combination of shotgun proteomics and RNA-seq. While the transcriptional profile of sgDHPS 3T3-L1 cells more closely mirrored WT differentiated versus undifferentiated cells, we observed early-on (i.e within the first 6-24 hours) striking differences in the translational control of differentiated sgDHPS 3T3-L1 versus differentiated WT 3T3-L1. It is important to note, though, that we have not excluded the possibility that loss of may DHPS lead to basal changes in mRNA and/or protein expression, as hypusination of EIF5A is likely critical elsewhere in general cellular function. It is clear, though, that in the presence of adipogenesis-inducing conditions that loss of DHPS specifically leads to early defects in translation but not transcription. One caveat to this data set, though, is that at later time points of adipogenesis (e.g. 24-96hrs) the accumulation of defects in translation will likely also lead to transcriptional defects, as many of the factors translated early on are transcription factors. Therefore, it would be difficult to distinguish between translational versus transcriptional control. Thus, we decided to restrict our search for transcriptional versus translational-dependent expression to the earliest time points in this experiment (6-24hr).

It is unclear if the failure to undergo MCE in the absence of hypusination is a consequence of loss of any single protein or instead the consequence of the sum of protein losses. The list of candidates regulated by EIF5A (as generated by our genome-wide CRISPR screen) contains both known and novel adipogenic regulators. However, our data suggest that even the known adipogenic regulators, such as BRD2 and KDM1A, are critical for adipogenesis because they are specifically required for MCE. Follow-up investigation will establish how these proteins function to promote MCE. Our shotgun proteomics data of differentiated sgDHPS versus WT 3T3-L1 also revealed that DHPS likely has roles in translational control beyond that of regulating the translation of proteins with stretches of multiple prolines. Of the 261 proteins that significantly increase in WT 3T3-L1 at any time point after induction with differentiation cocktail, 143 of these proteins either do not change or even show a significant decrease in sgDHPS cells upon differentiation cocktail treatment. Of these 143 proteins, though, only 33 had long stretches of prolines. This would suggest that the hypusination pathway has yet other roles to play in translational control beyond that of regulating the translation of proteins with long proline stretches.

### CRLs regulate lipogenesis

The screen identified that neddylation is strongly required for differentiation due to its requirement in MCE. In addition, NEDD8 had a significant positive LIPID score, arguing that neddylation inhibits lipid accumulation. Here, we show that neddylation of a CRL is required to prevent premature activation of insulin signaling and thus lipogenesis. We propose that this CRL is composed of CUL5 and ELOB/ELOC. Based on our proteomics analysis and previously published data (Hu et al., 2012), we further propose that IRS1 is a critical substrate of this CRL. We note that our proteomics analysis identified several other components of insulin signaling which are also stabilized as a consequence of neddylation inhibition, including PIK3R1, SRFBP1, PHIP, and IGFBP3. Moreover, PANTHER pathway analysis of differential protein abundance shows enrichment of pathways other than cell cycle and insulin. Further investigation is required to establish any of these additional proteins as CLR substrates.

Inhibition of neddylation promotes lipid accumulation in both preadipocytes and hepatocytes. Obesity is associated with an increased risk for hepatic steatosis and non-alcoholic fatty liver disease. Our data suggests that loss of NEDD8 and CRL function can drive this phenotype. Further investigation is needed to establish if dysregulation of this pathway contributes to human disease.

### Systems level analysis to define the adipogenesis and lipogenesis programs in new contexts

The list of strong genetic requirements for adipogenesis and lipogenesis that we discovered here facilitates many specific functional tests in preadipocytes and other cell types. The genome-wide comprehensive screen defines a core set of adipogenic subprograms for both known and new pathways causing specific metabolic changes throughout adipogenesis. This systems-level analysis is enabled by a global map of RNA and protein abundance through differentiation, coupled to knowledge of which gene products are critical. Specific subsets of these subprograms may be employed in other tissues that create fat deposits including bone marrow, muscle, liver, or specific fat depots. These subprograms include the translational and protein stability programs outlined here. Single cell profiles and shotgun proteomics in specific tissues may help pinpoint which of these adipogenic subprograms operate in specific subpopulations of preadipocytes and fat. New efforts can generalize from the preadipocyte requirements defined here to define how the adipogenesis program is molded to adapt fat tissue for specific needs in different physiological contexts.

## Acknowledgments

PKJ was supported by NIH grants 5R01GM114276, 5U01CA199216, 5UL1TR00108502 and the Stanford Department of Research and Baxter Laboratory. RET was supported by the Stanford Dean’s Fellowship and by the NIH 1F32GM142180-01A1. MCB was supported by NIH Director’s New Innovator Award (1DP2HD084069-01). KJS was supported by the NIH grants DK125260, DK111916, the Jacob Churg Foundation and the McCormick and Gabilan Award, and the Stanford Diabetes Research Center NIH grant P30DK116074. KIH was a Layton Family Fellow of the Damon Runyon Cancer Research Foundation (DRG-2210-14) and this project was supported by a ChEM-H Seed grant to postdocs at the interface to KIH and KH. CTJ was supported by an NIH training grant (TG2-01159). KH was supported by the Walter V. and Idun Berry award. AR was supported by NNF16OC0018642 from Novo Nordisk Foundation to the Stanford Bio-X program. RC was supported by the Cellular and Molecular Biology Training Grant (NIH 5 T32 GM007276). We thank N. Mooney for experimental support and members of the Jackson Laboratory for insightful discussions.

## Author Contributions

Conceptualization, RET, KIH, CTJ, MCB, and PKJ; Methodology, RET, KIH, CTJ, KH; Software, KH, AR and JD; Investigation, RET, KIH, CTJ, AR, RC, YZ, and ZJ; Resources, MCB, KJS, and PKJ; Writing – Original Draft, KIH and CTJ; Writing – Review & Editing, RET, KIH, CTJ, AR, and PKJ; Supervision, KIH and PKJ; Funding Acquisition, RET, KIH, KH, MCB and PKJ.

## Supplemental Information titles and legends

**Figure S1:**
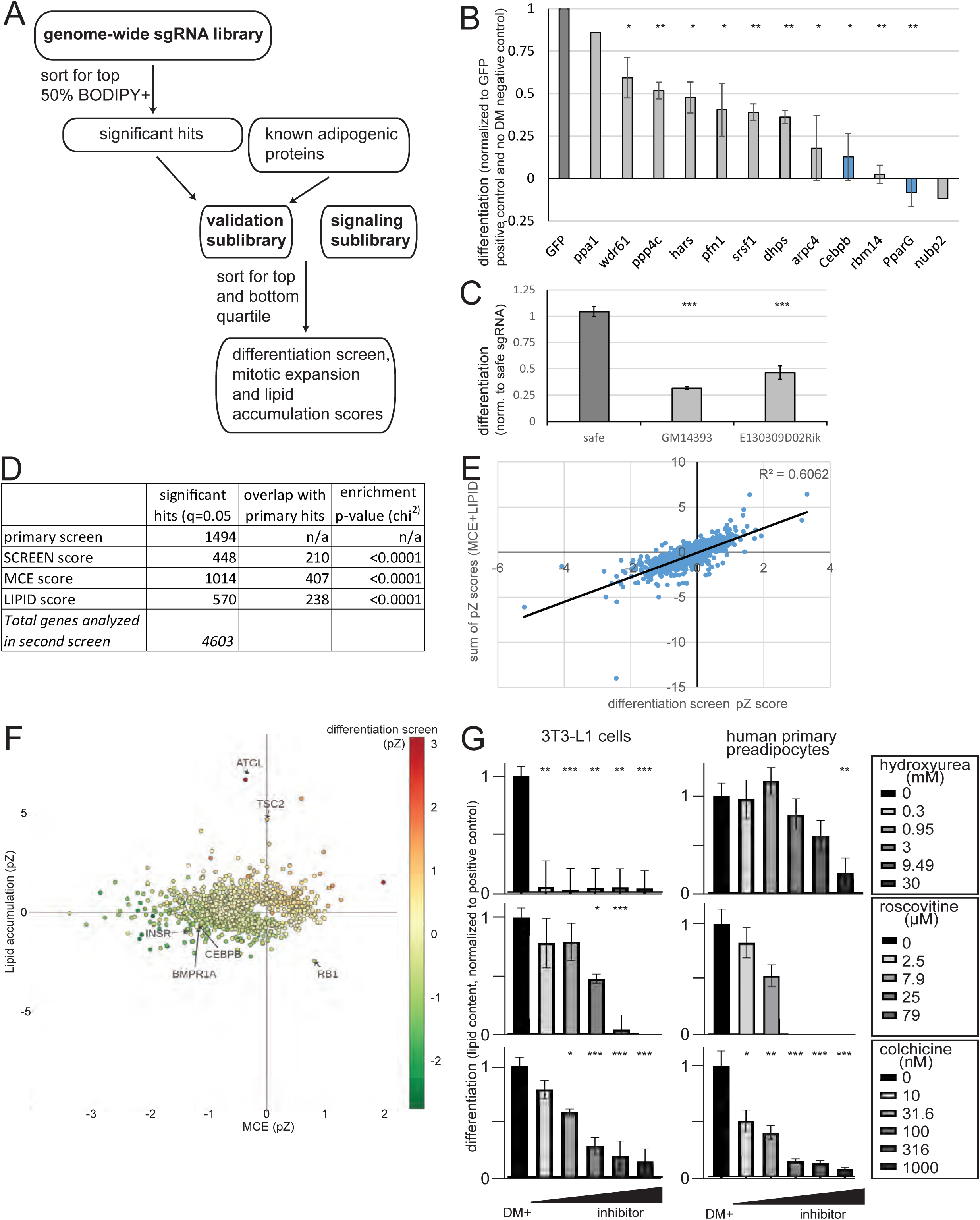
related to Figure 1. Genome-wide sgRNA screen of 3T3-L1 adipogenesis screen using a lipid-based sorting strategy. a. Workflow of primary and secondary screen. 3T3-L1 cells expressing CAS9 were transduced with a sgRNA library targeting all genes in the genome targeted with 10 distinct sgRNAs. The pool of knockout cells was differentiated in duplicate, treated with the BODIPY fluorescent dye, and the top 50% of differentiated cells were sorted as determined by green fluorescent intensity. Effect of each sgRNA was determined by comparing abundance relative to non-targeting control sgRNA in the Day 0 undifferentiated cell population versus Day 4 BODIPY-positive cell population. Significance was set at an FDR of 5%. Custom libraries of significant hits, any additional known adipogenic regulators based on literature search, and gene targets involved in signaling were generated. 3T3-L1 cells expressing CAS9 were transduced with the sgRNA sublibrary (10 sgRNAs pre gene). The pool of knockout cells was differentiated in duplicate, treated with the BODIPY fluorescent dye, and the top and bottom quartile of differentiated cells were sorted as determined by green fluorescent intensity. Effect of each sgRNA on differentiation, MCE, and lipid accumulation was determined by comparing relative sgRNA abundance between cell populations. b. 3T3-L1 preadipocytes lacking one of 10 genes identified in screen 1 were generated and differentiated. C/EBPB and PPARG were used as positive controls. Experiments were conducted in triplicates and bar graphs with error bars represent average of two independent experiments, each conducted in triplicates and normalized to its GFP positive and negative control. Error bar is ±SD. * p<0.05; ** p<0.01 c. 3T3-L1 preadipocytes lacking one of two uncharacterized genes, GM14393 and E130309D02Rik, that scored as significant in screen 1 were generated and differentiated. Experiments were conducted in triplicates and bar graphs with error bars represent average normalized to safe sgRNA control. Error bar is ±SD. *** p<0.001 d. The primary screen identified 1494 significant hits (FDR of 5%). A total of 4603 genes were tested in the secondary screen. This screen identified 448 hits important for differentiation, 1014 hits important for MCE, and 570 hits important for lipid accumulation (FDR of 5%). Hits identified in each category in the secondary screen were significantly enriched for hits identified in the primary screen as determined by a chi-squared test. This argues that the primary screen was successful in narrowing down the list of real adipogenic regulators. e. Scatter plot showing effect size in the differentiation screen versus the sum of the effect sizes obtained in the MCE and lipid accumulation screens. Only gene targets that scored as significant in at least one of the screens are plotted. More than half of the differentiation scores can be explained by the simple addition of the two components of adipogenesis, MCE and lipid accumulation. f. Scatter plot showing effect size in MCE screen versus lipid accumulation screen. Color gradient describes differentiation effect size. Only gene targets that scored as significant in at least one of the screens and with at least 12 sgRNA elements (out of 20 total in duplicate runs) recovered are plotted. Gene targets listed in Figure 1D are annotated.

**Figure S2: related to Figure 2.**
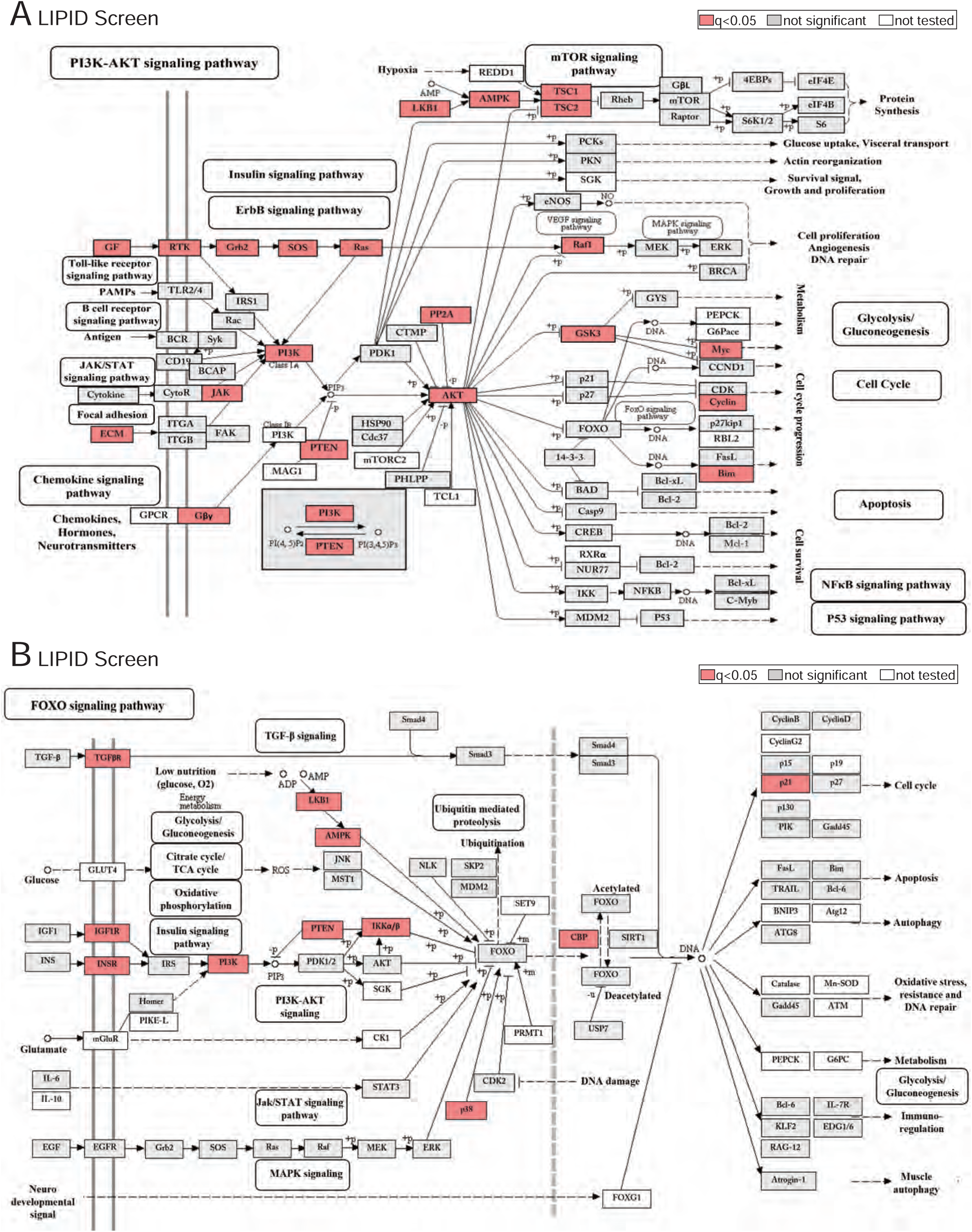
Adipogenesis screen results. a. KEGG pathway of PI3K-AKT signaling with gene targets that scored significantly in lipid accumulation screen denoted in red. b. KEGG pathway of FOXO signaling with gene targets that scored significantly in lipid accumulation screen denoted in red.

**Figure S3: related to Figure 3.**
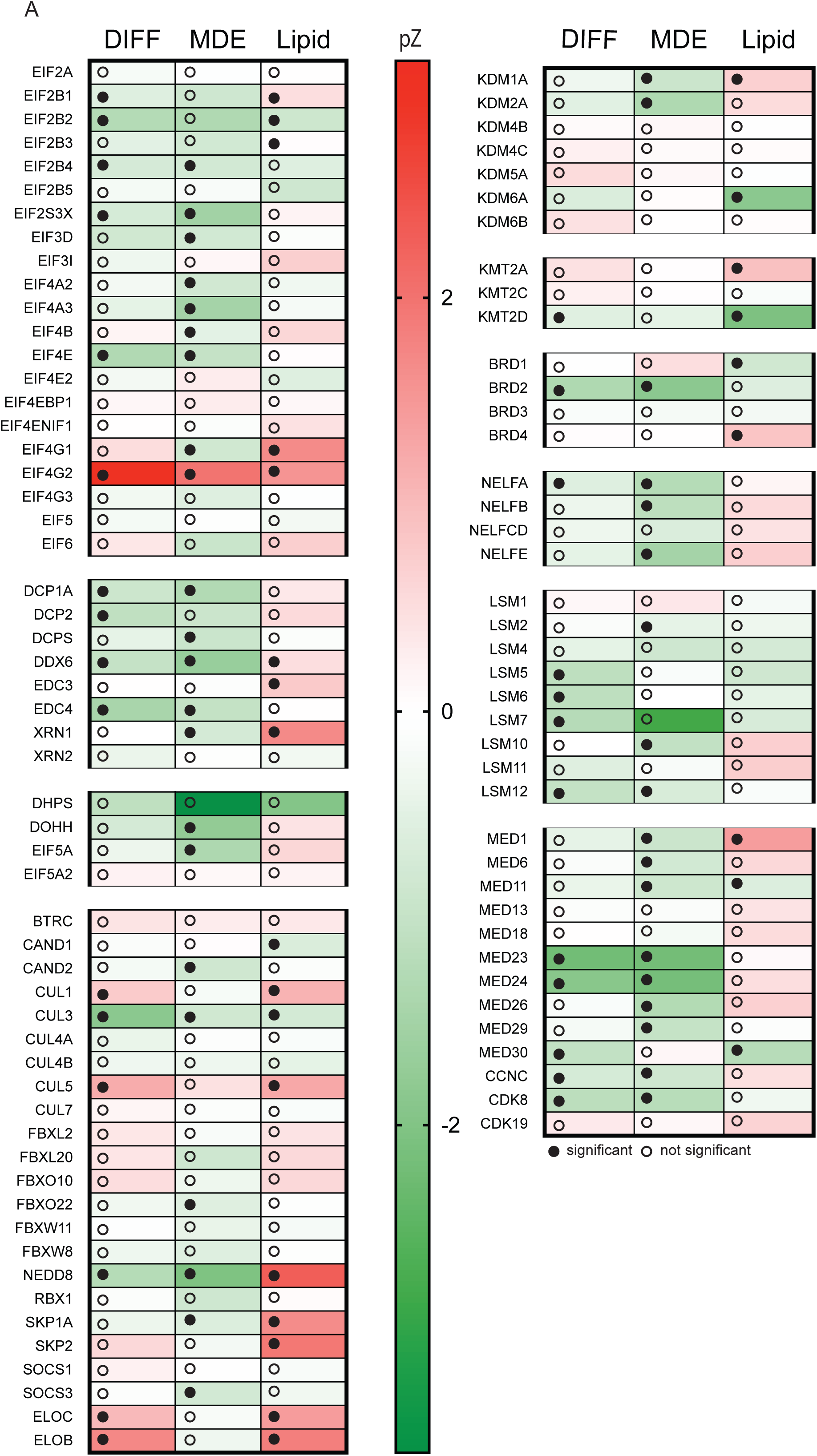
Heatmap of screen scores for genes in Figure 3. Heatmap displays phenotypic scores for complexes important for adipogenesis. Significance, qValue < 0.05, is denoted by filled circles.

**Figure S4: related to Figure 4.**
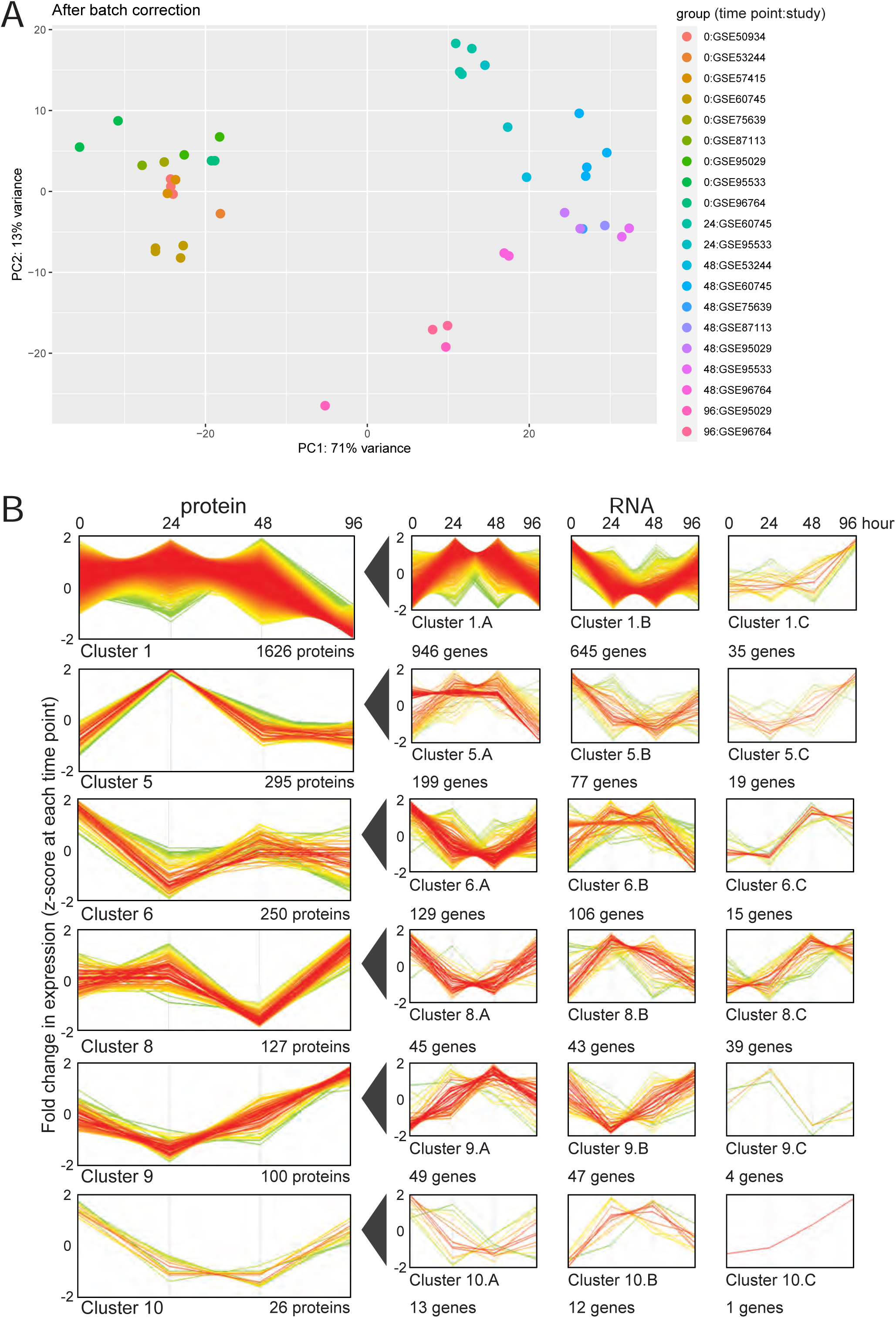
post-transcriptional regulatory mechanisms are critical to describe changes in expression during adipogenesis. a. PCA analysis of publicly available RNAseq data at 0, 24, 48, and 96 hour post-differentiation post-batch correction. RNA expression clusters by time point and independent of experimental source. b. Profile plot showing fold change in protein expression over time (after z-scoring for each time point) for six clusters as defined in Fig. 4C. RNA concentrations corresponding to the proteins in each cluster were analyzed and clustered hierarchically to yield three profile plots. Number of proteins/genes in each cluster is indicated below each cluster. Color of profile denotes distance from center, where red is zero and green is maximal change.

**Figure S5: related to Figure 5.**
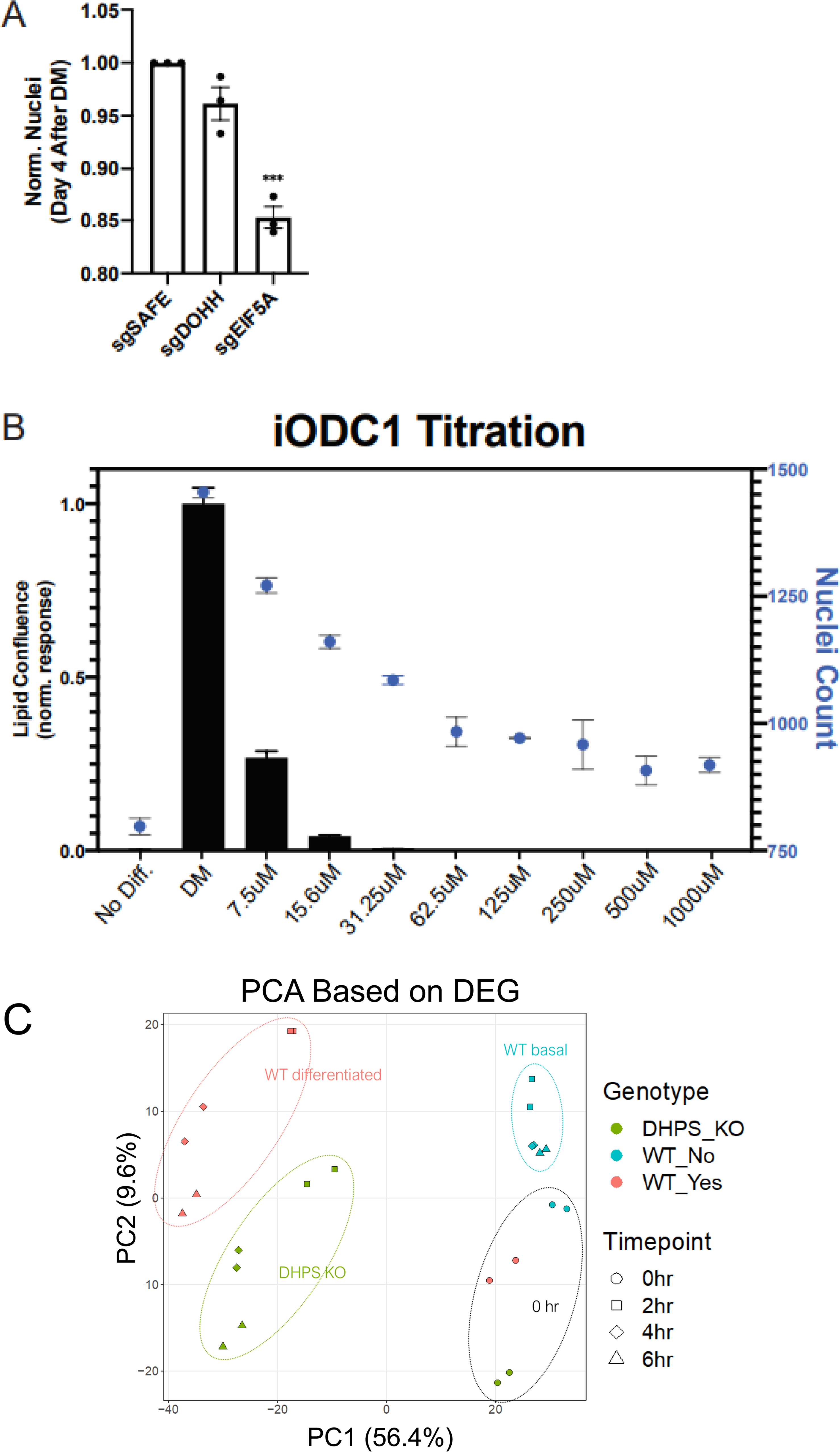
Inhibition or knockout of genes required for EIF5A hypusination lead to a loss of MCE in 3T3-L1 preadipocytes a. 3T3-L1 preadipocytes without DOHH or EIF5A have reduced Nuclei Count at 4 days post cocktail, consistent with a failure to undergo MCE in the absence of hypusination. Bar graphs represent normalized nuclei count ±SE *** p<0.001. **b.** 3T3-L1 preadipocytes were treated with 1mM aminoguanidine and the labeled concentrations of DMFO (iODC1) for two days prior to and after cocktail addition (4 days total). Lipid Confluence (black, left axis) and Nuclei Count (blue, right axis) were imaged and assessed 4 days after cocktail addition. Inhibition of ODC1 leads to a rapid loss of lipogenesis, and a nearly complete abrogation of MCE. Bar and scatter dot represent Lipid Confluence and Nuclei Count, respectively, ±SE.

**Figure S6: related to Figure 6.**
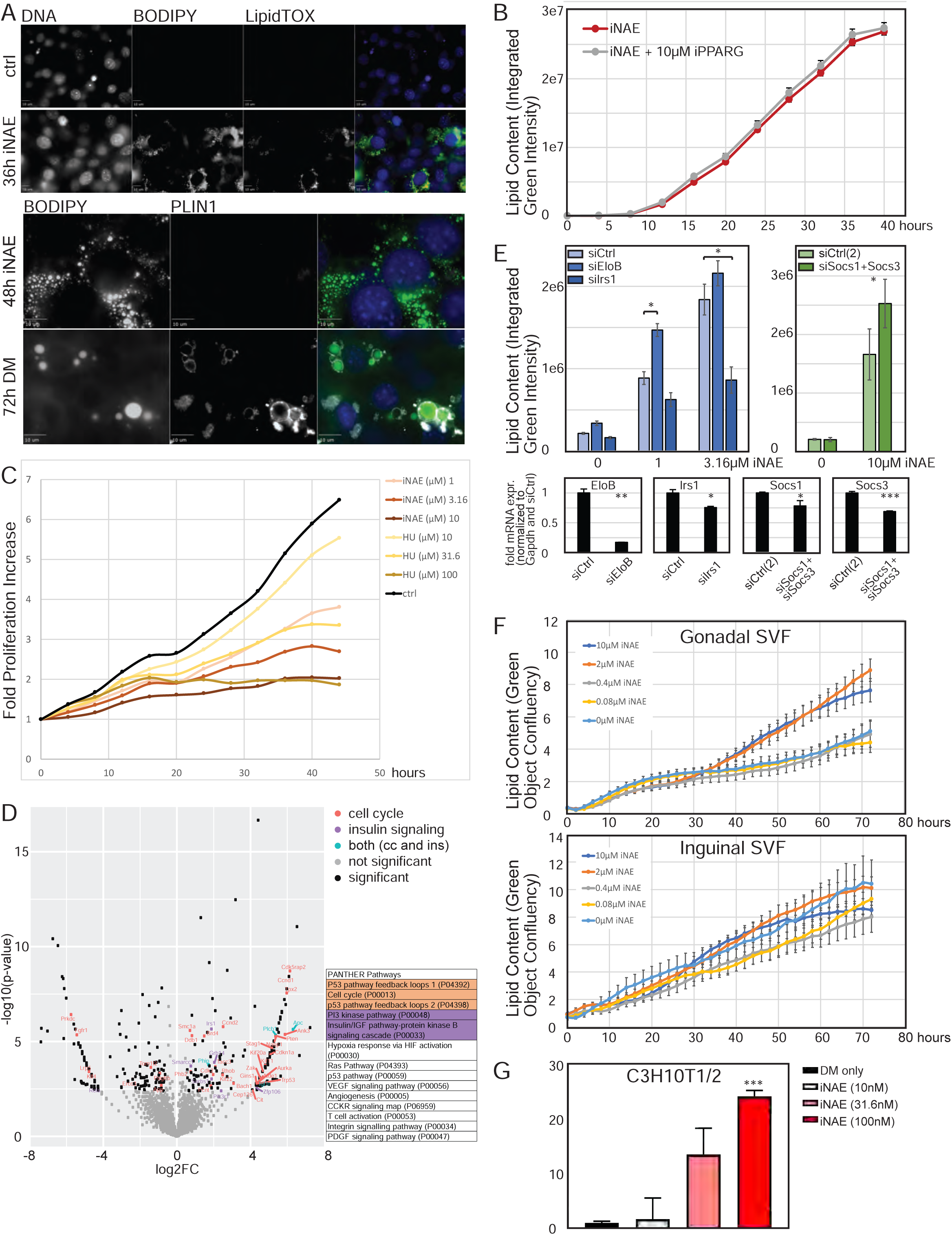
inhibition of neddylation promotes accumulation of lipids. a. 3T3-L1 preadipocytes treated for 36 hours with an inhibitor of neddylation accumulate lipid as determined by staining with the lipophilic fluorescent dyes BODIPY and LipidTox. This intracellular lipid appears fragmented and disorganized as determined by lack of PLIN staining after 48 hours of neddylation inhibition. As a positive control 3T3-L1 cells undergoing normal adipogenesis were stained with PLIN at 72 hours post differentiation media (DM) addition. Scale bar is 10µm. b. 3T3-L1 preadipocytes treated with an inhibitor of neddylation accumulate intracellular lipid even in the presence of a PPARγ inhibitor. Data points represent average of 3 independent experiments ±SE. c. 3T3-L1 preadipocytes treated with an inhibitor of neddylation or with an inhibitor of proliferation show decreased doubling time in a dose-dependent manner as measured by the confluency mask of the Essen Incucyte live-imaging system. d. Volcano plot of q-value versus log_2_ fold change of protein abundance in 3T3-L1 preadipocytes treated with vehicle versus inhibitor of neddylation for 14 hours as determined by shot-gun proteomics (left). Samples were run in triplicates. Panther pathway analysis of all protein with a log_2_ fold change greater than 1 (and identified in at least 3 out of 6 samples) shows an enrichment for proteins associated with cell cycle regulation and insulin signaling (right). Orange denotes cell cycle regulators, purple denotes components of insulin signaling, and teal describes hits associated with both the cell cycle and insulin signaling. e. Knockdown of EloB or Socs1 and Socs3 in 3T3-L1 preadipocytes promotes lipid accumulation, further potentiated by the addition of neddylation inhibitor. Knockdown of Irs1 in 3T3-L1 preadipocytes attenuates lipid accumulation, including in the presence of iNAE. Bar graphs are the average of 2 independent experiments ±SE. *Bottom*: quantification of mRNA expression upon knockdown of EloB, Irs1, or Socs1 and Socs3. Bar graphs are average of 3 technical replicates. * p<0.05; ** p<0.01; *** p<0.001; f. Inhibition of neddylation in primary gonadal preadipocytes promotes lipid accumulation in a dose-dependent manner as measured by BODIPY fluorescent intensity (top). Inhibition of neddylation in primary inguinal preadipocytes does not promote lipid accumulation (bottom). Data points represent average of 6 independent experiments for samples treated with iNAE and 3 independent experiments for control (0µM iNAE) ±SE. g. Inhibition of neddylation in C3H10T1/2 cells promotes lipid accumulation in a dose-dependent manner (48 hour treatment with iNAE). Data points represent average of 3 independent experiments ±SE.*** p<0.001;

**Figure S7:**
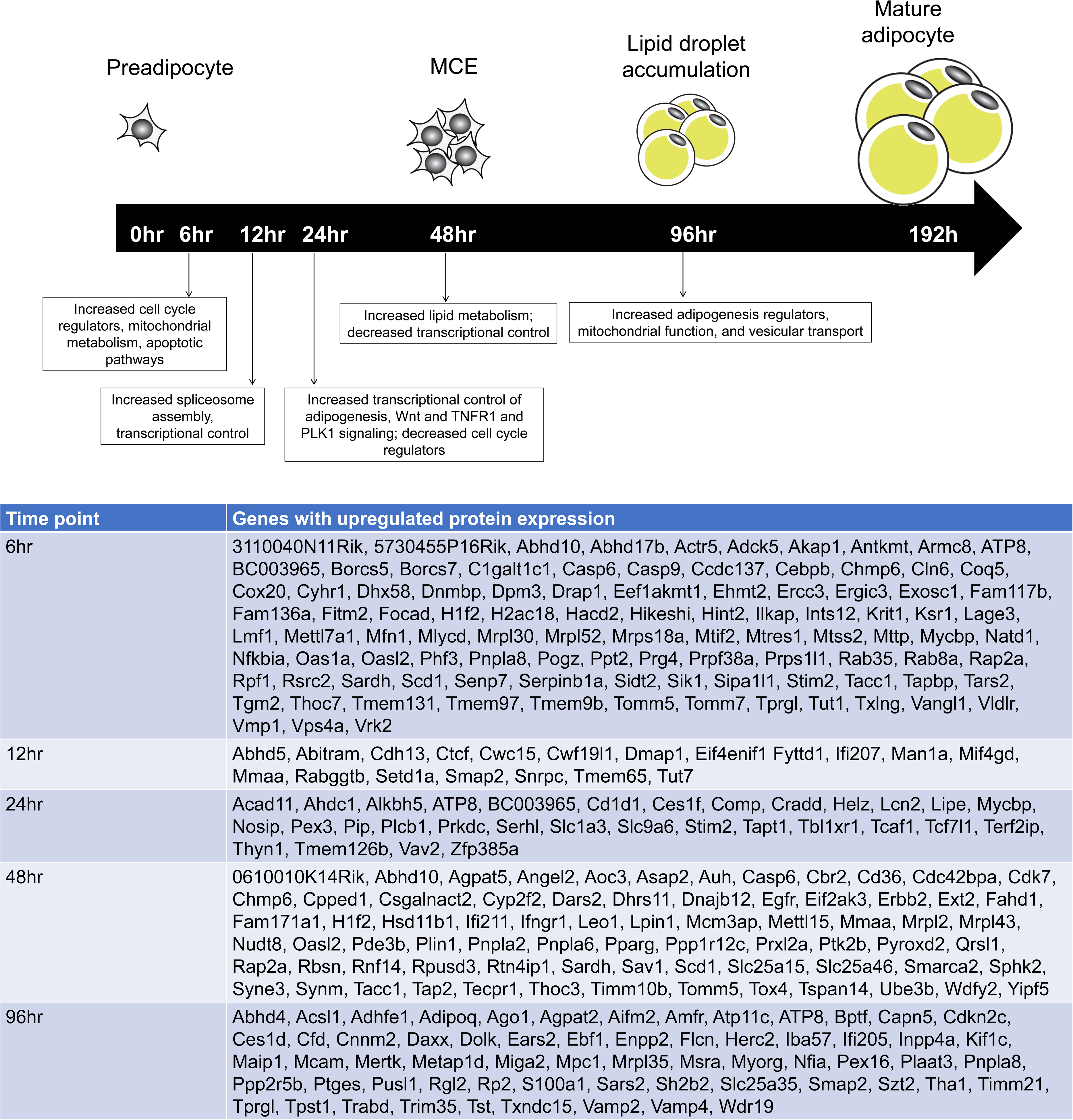
Detailed changes in transcriptional and translational control compared to Gene Ontology Mapping.

**Supplemental Table 1: Raw data of first screen Supplemental Table 2: Raw data of second screen**

**Supplemental Table 3: Protein expression during 3T3-L1 adipogenesis**

**Supplemental Table 4: Hierarchical clustering of changes in protein and RNA expression during 3T3-L1 adipogenesis**

**Supplemental Table 5: Neddylation inhibitor proteomics**

**Supplemental Table 6: Sublibraries**

**Supplemental Table 7: Primers used**

## Star Methods

### RESOURCE AVAILABILITY

#### Lead contact

Further information and requests for resources and reagents should be directed to and will be fulfilled by the Lead Contact, Peter Jackson.

#### Materials availability

All unique/stable reagents generated in this study are available from the Lead Contact with a completed Materials Transfer Agreement.

#### Data and code availability

The published article includes all datasets generated during this study. Code used to analyze datasets are available at https://github.com/biohank/CRISPR_screen_analysis

### EXPERIMENTAL MODEL AND SUBJECT DETAILS

#### *In vivo* animal studies

C57Bl/6J mice were purchased from Jackson Laboratory (000664). All mice were maintained at the Stanford animal care facility and all experiments were approved by the Administrative Panel on Laboratory Care at Stanford University. Animals were housed in groups of 5 adults per cage. Male mice between 6-8 weeks of age were used for isolation of murine primary preadipocytes.

#### Cell line models

3T3-L1 cells were cultured in DMEM medium containing 10% Bovine Calf Serum, 1% Pen/Strep, and 1% GlutaMAX and switched to DMEM containing 10% FBS, 1% Pen/Strep, and 1% GlutaMAX during adipogenesis.

Murine primary preadipocytes were isolated from inguinal or epididymal white adipose tissue from C57Bl/6J male mice using mouse protocols approved by the Institutional Animal Care and Use Committee (IACUC) at Stanford University. Primary preadipocytes were maintained and differentiated in DMEM medium containing 10% FBS, 1% Pen/Strep, and 1% GlutaMAX.

AML12 mouse hepatocytes were purchased from ATCC (#CRL-2254) and cultured with DMEM/F12 medium (Gibco) supplemented with 10% FBS, 10 µg/ml insulin, 5.5 µg/ml transferrin, 5 ng/ml selenium, 40 ng/ml dexamethasone, and 15 mM HEPES in a humidified atmosphere containing 5% CO_2_ at 37°C. AML12 hepatocytes were passaged once reached confluency.

### METHOD DETAILS

#### Plasmids

pMCB306 and pMCB302 (lentiviral vector, loxP-mU6-sgRNAs-puro resistance-EGFP-loxP and loxP-mU6-sgRNAs-puro resistance-mCherry-loxP) and p293 Cas9-BFP were gifts from Prof. Michael Bassik (Stanford University). pCMV-VSV-G and pCMV-dR8.2 dvpr were gifts from Bob Weinberg (Addgene plasmid #8454; http://n2t.net/addgene:8454 ; RRID:Addgene_8454 and #8455; ; http://n2t.net/addgene:8455 ; RRID:Addgene_8455) (Stewart et al., 2003). pHCMV-EcoEnv was a gift from Miguel Sena-Esteves (Addgene plasmid # 15802; http://n2t.net/addgene:15802; RRID:Addgene_15802) (Sena-Esteves et al., 2004). Lentiviral vectors containing sgRNA were generated by ligating annealed sgRNA oligonucleotides into pMCB306 vector digested with BstXI and BlpI restriction enzymes.

#### Cell Line generation

Lentiviral vectors carrying the gene of interest were co-transfected with pCMV-VSV-G or pHCMV-EcoEnv and pCMV-dR8.2 dvpr into 293T cells using Fugene6 (Promega). Media was replaced after 24h and virus was harvested 48 and 72h post-transfection. Virus was filtered with a 0.45μm OVDF filter (Millipore) or freeze-thawed and subjected to centrifugation at 3000rpm for 10min. Cell lines were infected with virus in 10μg/ml polybrene (Millipore). Media was replaced after 24h and infected cells were isolated either via antibiotics selection or FACS sorting after 48-72h post-infection.

3T3-L1 cells expressing Cas9-BFP were generated by infection of virus harvested from 293T cells transfected with p293 Cas9-BFP, pCMV-VSV-G and pCMV-dR8.2 dvpr. 3T3-L1 Cas9-BFP cells were sorted for BFP positivity. To generate Crispr/Cas9 knockout cells, 3T3-L1 Cas9-BFP cells were infected with lentivirus containing the sgRNA of interest. Primers listed in Table S7.

3T3-L1 sgRNA pools for screening were generated as follows: sgRNA sublibraries were cloned from oligonucleotide pools (Agilent) as previously described (Morgens et al., 2017). Genome-wide sgRNA library was synthesized in 20 sublibraries with 10 different sgRNAs per gene and we generated 2 additional sublibraries for second screen (Table S6). Oligonucleotides in each sublibrary were amplified with a sublibrary specific library adaptor protein, digested with BstXI and BlpI restriction enzymes, and ligated into pMCB320 vector. 10μg of each sublibrary, 7μg of pCMV-dR8.2 dvpr and 3μ of pHCMV-EcoEnv was co-transfected using Fugene6 and OptiMem onto one 15cm plate of 293T cells. Media was replaced after 24h and 25ml of viral supernatant was collected each at 48 hours and 72 hours post-transfection and frozen. For each sublibrary, 1.5e6 3T3-L1 Cas9-BFP cells were plated into 1050 cm^2^, smooth PETG roller bottles (Fisher, 12-575-120). 96 hours post-plating, 50ml of viral supernatant of each sublibrary was thawed, spun for 10min at 300rpm and added to each roller bottle in a total of 200ml media volume containing 10μg/ml polybrene. After 24 hours, cells were trypsinized and passaged into a 1800 cm^2^, smooth roller bottles (Fisher, 12-565-529). After 24 hours, cells were selected in 2 μg/ml puromycin. Cells were grown for 2 weeks, maintained at any time at 1000x the number of sgRNA elements present in the sublibrary (∼1000 genes per sublibrary, resulting in ∼10,000 sgRNA elements per sublibrary, requiring maintenance at ∼10 million cells per sublibrary), and then frozen as separate sublibrary pools in multiple aliquots of 10^7^ cells. 1 aliquot of each sublibrary was subsequently combined for each experimental replicate and time-point.

#### *In vitro* Adipogenesis

3T3-L1 cells were grown to confluency in DMEM containing 10% Bovine Calf Serum, followed by another 2 days at confluency in DMEM containing 10% Bovine Calf Serum. Adipogenesis was then induced using DMEM containing 10% FBS and differentiation cocktail consisting of 1μg/ml insulin, 1μM Dex, and 0.5mM IBMX. After 2 days of differentiation cocktail, media was changed to DMEM containing 10% FBS and 1μg/ml insulin. Maintenance media was changed every 2-3 days for a total differentiation time of 4-8 days. Where indicated, cells were treated with inhibitors to enzymes in the hypusination pathway for a total of four days, starting 2 days prior to addition of differentiation cocktail and ending on Day 2 of differentiation. Cells treated with hypusination inhibitors GC7 and DFMO were supplemented with 1mM Aminoguanidine.

Mouse primary preadipocytes were grown to confluency in DMEM containing 10% FBS. Adipogenesis was induced using DMEM containing 10% FBS and differentiation cocktail consisting of insulin, dexamethasone, IBMX and/or 10μM NAE inhibitor MLN4924 (Selleckchem, S7109, Pevonedistat). After 3 days of differentiation cocktail, media was changed to DMEM containing 10% FBS, 1μg/ml insulin and 10μM NAE inhibitor MLN4924. Maintenance media was changed every 2-3 days.

#### Functional genomics screen

The screen was conducted in duplicate. For each condition (Day 0 bulk, Day 4 bulk, Day 4 for sorting), 1 aliquot of 10^7^ cells for of each 3T3-L1 Cas9 sublibrary pool was combined in 30ml of fresh media and centrifuged for 5min at 1300g. Screen was performed in duplicates/condition. Pellets were resuspended and filtered using 100µm cell filter. Cell pools were then plated in multiple 1800 cm^2^, smooth roller bottles such that each roller bottle contained equivalent numbers of cells (5×10^7^ cells) from only 1 condition and replicate. Cells were grown to confluence, followed by media change. Cells were kept at confluency for 48 hours.

Day 0 bulk samples were collected by trypsinization and frozen in media supplemented with 10% DMSO. For remaining samples, adipogenesis was induced using DMEM containing 10% FBS and differentiation cocktail consisting of 1μg/ml insulin, 1μM Dex, and 0.5mM IBMX. After 48 hours, the medium was changed to DMEM containing 10% FBS and 1μg/ml insulin. After 48 hours, Day 4 bulk samples were collected by trypsinization and frozen in media supplemented with 10% DMSO. To the remaining samples, BODIPY 493/503 (ThermoFisher, D-3922) was added to media for a final concentration of 5µM for 1 hour at 37°C (incubator). Cells were collected by trypsinization, combined into each duplicate and filtered using 100µm cell filter. Cells were washed with PBS and pellets were resuspended in PBS containing 1% serum.

Samples were gated for mCherry+ and FITC as indicated in the figure legend (top 50% for first screen, top and bottom 20% for second screen) on a FACSAria II (BD) instrument. Sorted populations were pelleted and frozen in media supplemented with 10% DMSO. DNA was isolated with a QiaAMP DNA Blood Maxi Kit (Qiagen) concurrently for all samples. Genomic DNA was then amplified with Herculase II polymerase (Agilent) as previously described (Deans et al., 2016). Briefly, outer primers were used to amplify the sgRNA cassette and inner primers were used to amplify the PCR product and to introduce Illumina sequencing barcodes and adapters (Table S7). PCR products were gel-extracted, quantified using dsDNA HS kit (Thermo Fisher), and pooled for deep sequencing using NextSeq 500 sequencer with high-output v2 kits (Ilumina). Sequencing was performed using a custom primer against the sgRNA protospacer.

Phenotypes from CRISPR screens were analyzed as previously described (Han et al., 2020). Briefly, log2 fold enrichments (log2e) of sgRNAs were calculated between undifferentiated and differentiated 3T3-L1 samples and subsequently normalized to the median log2e of negative control sgRNAs (Morgens et al., 2017) in libraries. Log2e of all sgRNAs were then divided by the standard deviation of negative control sgRNAs to yield phenotype Z scores (pZ) of sgRNAs. A median pZ score of ten sgRNAs for a given gene was measured and used as the phenotype of the gene. The probability that the distribution of all ten sgRNAs targeting a given gene was significantly different from that of negative control sgRNAs was measured using Mann-Whitney U (MWU) test to calculate statistical significance of gene phenotypes. Lastly, to correct for multiple hypothesis testing, we calculated the adjusted p values (q values) based either on the genome-wide distribution of p values (Storey and Tibshirani, 2003) or on the Benjamini– Hochberg procedure implemented in statsmodels, a python statistical module (https://www.statsmodels.org/stable/index.html). pZ scores of sgRNAs from two experimental replicates were pooled together to calculate combined phenotypes of genes. Scripts used to analyze the data were published previously (Han et al., 2020) and are available at https://github.com/biohank/CRISPR_screen_analysis.

To overlap screen hits with GWAS, GWAS summary statistics were obtained from the publicly available GIANT consortium’s download site: http://portals.broadinstitute.org/collaboration/giant/images/f/f0/All_ancestries_SNP_gwas_mc_merge_nogc.tbl.uniq.gz.

#### Flow cytometry analysis

Cells were treated with 5µM BODIPY 493/503 for 1 hour at 37°C and collected by trypsinization. Cells were washed twice with PBS and pellets were resuspended in PBS containing 1% serum. Samples were analyzed for FITC on a FACSCalibur (BD) and data was analyzed by FlowJo (TreeStar).

#### KEGG pathway and Gene Ontology enrichment analysis

Kyoto Encyclopedia of Genes and Genomes (KEGG) pathway analysis was conducted to identify the biological role of the hits from our screen (PMID: 30321428, PMID: 10592173, PMID: 27899662). All the genes recovered in the second screen were used as background for the enrichment analysis. KEGG pathway analysis revealed that hits from MCE screen were enriched for cell cycle activity, the hits from LIPID screen were mainly involved in TCA cycle and oxidative phosphorylation, and the hits from SCREEN screen were enriched for insulin signaling, PI3K-AKT, mTOR, and FOXO pathways. Genes visualized in white were not tested in the second screen, genes visualized in grey were tested but did not score as significant hits in MCE, LIPID, or SCREEN, and genes visualized in light red were tested and scored as significant (adjusted P<0.05) in one or more of the analyses (MCE, LIPID, SCREEN).

Gene Ontology enrichment for biological processes was performed on all significant SCREEN hits (q<0.05) with a background of all genes analyzed in the second screen. Panther Pathway Analysis was performed on hits from the neddylation inhibitor proteomics dataset. 3T3-L1 cells in triplicates were treated with 10µM NAE inhibitor MLN4924 for 14 hours. Hits were defined as having a peptide count in at least 3 out of the 6 samples and at least a 2 fold increase in expression in the presence of the inhibitor.

#### Combining public RNA expression data sets

Publicly available RNAseq data sets (Ahmed and Kim, 2019) were analyzed. Data sets used are available: http://bioconductor.org/packages/release/data/experiment/vignettes/curatedAdipoRNA/inst/doc/curatedadiporna_vignette.html

Briefly, low mean ‘per base sequence quality’ samples (< 34) were filtered out, and samples for 4 time points (0, 24, 28 and 96 hrs) were picked for further analysis. Low expression genes that did not meet the criteria of (count > 10 in at least 2 samples) were filtered out. The resulting dataset was analyzed by DESeq2 using time and study name as variables. Genes with log2-fold change > 1 and adjusted p-value < 0,05 were considered significant.

#### Proteomics analysis

3T3-L1 cells were cultured and differentiated as described above. Where noted, samples were treated with 10µM NAE inhibitor MLN4924 for 14 hours. At described time points, cells were washed twice with PBS and then collected using a cell scraper in PBS. Samples were centrifuged and pellets were flash-frozen in liquid nitrogen. Pellets were thawed and resuspended in 500µl freshly made Cell Lysis Buffer (50mM HEPES pH 8.5, 75mM NaCl, 8M Urea, and protease and phosphatase inhibitors (ThermoFischer, A32961). Cells were lysed using QIAshredder columns (Qiagen, 79656) at 5000rpm for 1 min at 4°C and clarified by centrifugation at 15,000rpm for 10min at 4°C. Protein concentration of supernatant was quantified using a BCA protein assay kit (ThermoFischer, 23225). 20µg of each lysate were reduce using 5mM TCEP (ThermoFischer, PG82080) for 25min at room temperature and alkylated using 10mM CAA (freshly made, Sigma, C0267) for 30min at room temperature in the dark, followed by quenching using 15mM DTT for 15min at room temperature in the dark.

Samples were precipitated by adding lysis buffer to final volume of 100µl, adding 100µl methanol and vortexing, 50ul chloroform and vortexing, and 100µl of water (for UHPLC; Sigma, 900682), followed by centrifugation at 14,000*g* for 3 min at 4°C. Methanol was removed followed by speed-vac to dryness. Samples were resuspended in 50µl of 200mM HEPES pH 8.5 for a final protein concentration of 0.4µg/µl. Samples were digested overnight by adding 0.2µg LysC and Trypsin (Promega, V5073; 10x stock was supplemented with 1% ProteaseMax (Promega, V2072) and Ammonium bicarbonate) and shaking at 250rpm at 37°C. Samples were acidified in TFA (ThermoFischer, L06374-AC) to reach pH 2 to stop digestion. Samples were cleaned up using C18 stagetips. Briefly, 5 layers of C18 (Empore, 320907D) were punched out with 0.9mm needle and packed into a 200µl pipette tip, resin was activated with 50µl methanol, followed by centrifugation at 2000g for 2min at 4°C. Stagetip was washed three times with 150 µl 5% acetonitrile (ACN; Sigma, 34998) 0.5% TFA, centrifuge 2000g for 2min at 4°C. Samples were bound by adding to C18 column and centrifuging at 500g for 5min at 4°C. Samples were washed twice with 150 µl 5% ACN 0.5% TFA and centrifugation at 2000g for 2min at 4°C. Samples were eluted with 25 µl 50% ACN 0.1% Formic Acid (FA; ThermoFischer A117-50) and centrifugation at 2000g for 2min at 4°C. Elution was repeated two more times followed by speed-vac. Peptides were resuspended in 20ul Solution A (2% CAN, 0.1% FA in water for UHPLC) and quantified using Quantitative Colorimetric Peptide Assay (ThermoFischer, 23275).

Samples were diluted to 100ng/µl and 200ng of sample was analyzed using the timsTOF Pro (Bruker Daltonics), an ion-mobility spectrometry quadrupole time of flight mass spectrometer. Specifically, a nanoElute (Bruker Daltonics) high pressure nanoflow system was connected to the timsTOF Pro. Peptides were delivered to the sample trap (5 cm x 300µm i.d., ThermoFisher Scientific, 160454) in line with a reversed phase analytical columns (25 cm x 75 μm i.d., IonOpticks, AUR2-25075C17A-CSI) with pulled emitter tips. Liquid chromatography was performed at 45 °C and peptides were separated on the analytical column using a 120 min gradient (solvent A: 2% ACN, 0.1% FA; solvent B: 0.1% FA, in ACN) at a flow rate of 400 nl/min. A linear gradient from 2-15 % B was applied for 60 min, followed by 15-23 % B for the next 30 min and followed by a step to 35% B for 10 min and a step to 80% B for 10 min followed by 10 min of washing at 80% B. The timsTOF Pro was operated in PASEF mode with the following settings: Mass Range 100 to 1700m/z, 1/K0 Start 0.6 V·s/cm2, End 1.6 V·s/cm2, Ramp time 100ms, Lock Duty Cycle to 100%, Capillary Voltage 1600V, Dry Gas 3 l/min, Dry Temp 200°C, PASEF settings: 10 MS/MS, charge range 0-5, active exclusion for 0.4 min, Scheduling Target intensity 20000, Intensity threshold 2500, CID collision energy 59eV.

Raw data files from Bruker timsTOF Pro were analyzed using Byonic software (Protein Metrics, Inc) to identify peptides and proteins with the following settings: precursor mass tolerance 20 ppm, fragment mass tolerance: 40 ppm using QTOF/HCD fragmentation type. Fully specific tryptic digest with maximum 2 missed cleavages were specified. The following modifications were searched for the differentiation time course: carbamidomethyl (fixed), Met oxidation, Ser/Thr/Tyr phosphorylation and Asn/Gln deamidation (common), N-term acetylation and Gln/Glu to pyro-Glu conversion (rare) with total common max = 2 and total rare max = 1. The following modifications were searched for the neddylation inhibition experiment: carbamidomethyl (fixed), Met oxidation, Ser/Thr/Tyr phosphorylation and Asn/Gln deamidation (common), N-term acetylation and GlyGly (for ubiquitin modification, neddylation, …) and ValGly (Ufm1 modification) modifications on Lys (rare) with total common max = 2 and total rare max = 1. We used a mouse refseq proteome library (June 2018) supplemented with common contaminants. To estimate false discovery rates, we applied a target-decoy library strategy using reverse sequences with FDR set at 1%. For differential expression analysis we used the msmsTests package.

#### Heatmap and Clustering

The log_2_ values of protein ratios were normalized by z-normalization among different time points. The z-normalized proteins were imported in Perseus (PMID: 27348712) and hierarchical clustering analysis was performed for proteins (rows). K-means clustering of all quantified proteins across the timespan resulted in 10 clusters. For clustering, we adopted an average linkage method, and also the Euclidean distance to compute pairwise distances. The profile plots of ten clusters with distinct behavior were generated in Perseus with the y-axis representing the z-score and the x-axis representing the time point. The colors represent the distance from the center with red being closer to the center of the trend and green being more far from the center.

Each of the ten protein clusters was categorized in 3 sub-clusters based on the regulation pattern of their corresponding RNAs.

#### qRT-PCR

3T3-L1 were harvested using 5mM EDTA in PBS, and the cells were lysed to harvest RNA using the RNeasy Lipid Tissue Kit (QIAGEN). cDNA was synthesized using M-MLV Reverse Transcriptase (Invitrogen, 28025-013). Quantitative real time PCR was performed using TaqMan Probes (Invitrogen) and the TaqMan Gene Expression Master Mix (Applied Biosystems, 4369016) in 96-well MicroAmp Optical reaction plates (Applied Biosystems, N8010560). Expression levels were normalized to the average expression of the housekeeping gene NoNo. The following probes were used All experiments were performed in technical triplicate on a Quant Studio 6 real-time PCR machine (Applied Biosystems). Fold changes were calculated using the 2-ΔΔCT method.

#### RNA-seq

As described above for qPCR, 3T3-L1

#### Sample preparation and immunoblot

Cells were lysed in 1x LDS buffer containing DTT and incubated at 95C for 5min. Proteins were separated using NuPage Novex 4-12% Bis-Tris protein gels (Thermo Fisher Scientific, WG1402BOX) in NuPage MOPS SDS running buffer (50 mM MOPS, 50 mM Tris Base, 0.1% SDS, 1 mM EDTA, pH 7.7), followed by transfer onto nitrocellulose membranes (Life Technologies, LC2001) in Towbin Buffer (25 mM Tris, 192 mM glycine, pH 8.3) containing 10% methanol. Membranes were blocked in LI-COR Odyssey Blocking Buffer (LI-COR, NC9232238) for 30 min at room temperature, followed by incubation with primary antibody in blocking buffer for at least 1h at room temperature. The membrane was washed 3 times for 10min in TBST buffer (20 mM Tris, 150 mM NaCl, 0.1% Tween 20, pH 7.5) at room temperature, incubated with secondary IRDye antibodies (LI-COR) in blocking buffer for 30 min at room temperature, and then washed 3 times for 10min in TBST buffer. Membranes were scanned on an Odyssey CLx Imaging System (LI-COR), with protein detection at 680 and 800 nm. Antibodies were used as follows: pIRS1 (Cell signaling 2381, 3203, 5610, 1:1000), pAKT (Cell signaling, 4051, 1:1000), Tubulin (Sigma, 9026, 1:5000).

#### Immunofluorescence and Lipid staining

Cells were grown on acid washed 12mm round coverslips and fixed with 4% paraformaldehyde (433689M, AlfaAesar) in PBS at room temperature for 10min. Samples were blocked with 5% normal donkey serum (017-000-121, Jackson ImmunoResearch) in IF buffer (3% BSA and 0.1% NP-40 in PBS) at room temperature for 30min. Samples were incubated with primary antibody in IF buffer at room temperature for 1h, followed by 5 washes with IF buffer. Samples were incubated with fluorescent-labeled secondary antibody at room temperature for 30min, followed by a 5 min incubation with 4’,6-diamidino-2-phenylindole (DAPI) in PBS at room temperature for 5min and 5 washes with IF buffer. Coverslips were mounted with Fluoromount-G (0100-01, SouthernBiotech) onto glass slides followed by image acquisition. Antibodies were used as follows: Anti-Perilipin-1 antibody (ab61682, 1:100).

To confirm fatty acid composition upon treatment with NAE inhibitor MLN4924, 3T3-L1 cells were plated onto coverslips and grown to confluency. Cells were treated with DMSO control or NAE inhibitor MLN4924 as well as methanol control or 10µM BODIPY C12-fatty acid (Invitrogen, D3822) for 32 hours. Samples were fixed in 4% paraformaldehyde in PBS at room temperature for 10min and stained with DAPI in PBS at room temperature for 5min. Coverslips were mounted with Fluoromount-G onto glass slides followed by image acquisition.

To stain lipids using BODIPY or LipidTOX, cells on coverslips were fixed in 4% paraformaldehyde in PBS at room temperature for 10min and stained with LipidTOX Deep Red Neutral Lipid Stain (ThermoFisher, H34477, 1:200), BODIPY 493/503 (ThermoFisher, D-3922, 20uM), and 1µg/ml DAPI in PBS at room temperature for 30min. Coverslips were mounted with Fluoromount-G onto glass slides followed by image acquisition.

#### Epi-fluorescence and confocal imaging

Images were acquired on an Everest deconvolution workstation (Intelligent Imaging Innovations) equipped with a Zeiss AxioImager Z1 microscope and a CoolSnapHQ cooled CCD camera (Roper Scientific) and a 40x NA1.3 Plan-Apochromat objective lens (420762-9800, Zeiss) was used.

Confocal images were acquired on a Marianas spinning disk confocal (SDC) microscopy (Intelligent Imaging Innovations).

#### Isolation of primary preadipocytes

Inguinal or epididymal white adipose tissue was removed and minced. Minced tissue was incubated in Collagenase Buffer (3,000 U/ml type II collagenase powder (Sigma, C6885), 100 U/ml DNase (Worthington, LS006344), 1 mg/ml poloxamer 188 (P-188) (Sigma P5556), 1 mg/ml BSA, 20 mM HEPES buffer, and 1 mM CaCl_2_ in Medium 199 with Earle’s salts (Sigma, M4530)) for 10min at 37C, followed by 20min shaking (250rpm) at 37C. Digested samples were strained through a 100μm filter and diluted 1:1 in cell suspension buffer (2% FBS, 1 mg/ml P-188, and 1% pen/strep in PBS), followed by centrifugation (1300rpm, 5min).

For isolation of SVF cells depleted for RBCs and WBCs, the cell pellet was resuspended in cell suspension buffer and layered onto Histopaq-1077 (Sigma, 10771), followed by 20min RT centrifugation at 2000rpm. The cell layer was removed and centrifuged (5min, 1300rpm), and cell pellet was resuspended in cell suspension buffer containing anti-CD45 microbeads (Miltenyi Biotec, 130-052-301) and anti-TER119 microbeads (Miltenyi Biotec, 130-049-901) for 30min at 4C, followed by MACS depletion using an LD column. Flow-through was collected and centrifuged (1300rpm, 5min). Cell pellet containing primary preadipocytes were resuspended in DMEM containing 10% FBS, 1% Pen/Strep, and 1% GlutaMAX.

#### Oil Red O staining and quantification

Cells are fixed in 4% PFA/PBS for 10min at room temperature, followed by 3 rinses in PBS. Samples were incubated in 60% isopropanol for 5min at room temperature and then allowed to dry completely. Samples are then incubated in freshly diluted 60% Oil Red O staining solution in water (stock is 0.5% Oil Red O (Sigma, 00625) in isopropanol) for 20min at room temperature, followed by 3 rinses in water. Samples were allowed to dry completely and imaged. To quantify, Oil Red O was extracted by incubating dried samples stained on the same day in 100% isopropanol for 5min at room temperature and absorbance was measured at 510nm.

#### Live Cell Imaging Analysis for lipid accumulation and proliferation

Live imaging for kinetics quantification of adipogenesis and for proliferation assay was performed using the IncuCyte Live Cell Analysis Imaging System (Essen Bioscience) with images acquired every 2h using the 10x objective. 3T3-L1 cells were grown in the presence of indicated concentrations of NAE inhibitor MLN4924, proteasome inhibitor MG132, and DNA replication inhibitor hydroxyurea (HU). Proliferation was assessed using a confluency mask generated by the IncuCyte Zoom Analysis Software using phase images. For the kinetic quantification of adipogenesis, 3T3-L1 cells were differentiated as described above or supplemented with inhibitors as described above and supplemented with 2µM BODIPY 493/503. Green fluorescence images were acquired every 2h using default IncuCyte setting. Total green fluorescence intensity was determined from a green fluorescent mask generated by the IncuCyte Zoom Analysis Software.

#### Glucose uptake assay

3T3-L1 cells were plated in a 96 well optical bottom plate and grown to confluency. Cells were treated with DMSO control or 10µM NAE inhibitor MLN4924 for indicated time periods in the presence of 100µM 2-NBDG (Cayman, 11046) or ethanol control. Cells were imaged using the IncuCyte Live Cell Analysis Imaging System (Essen Bioscience) after 2 PBS washes using default IncuCyte settings to quantify total green fluorescence intensity from a green fluorescent mask generated by the IncuCyte Zoom Analysis Software.

#### Lipogenesis assay

AML12 hepatocytes were washed with warm PBS twice and starved in serum-free DMEM/F12 overnight. Indicated concentrations of insulin with or without NAE inhibitor MLN4924 was added at the same time. After 24 hours incubation, a mixture of 10 µM cold acetate and 2 µCi 3H-Acetate (#NET003H005MC, PerkinElmer) was added to each well and cells were incubated for another 4 hr. Cells were washed with PBS twice and lysed using 0.1 N hydrogen chloride. Lipids were extracted by 2:1 chloroform-methanol (v/v). After 10 min centrifugation at 3000 x g, lower phase was transferred to a scintillation vial, and 3H activity was measured by liquid scintillation counting.

### QUANTIFICATION AND STATISTICAL ANALYSIS

Statistical parameters including the statistical test used, exact value of n, what n represents, and the distribution and deviation are reported in the figures and corresponding figure legends. Most data are represented as the mean ± standard deviation and the p-value was determined using two-tailed Student’s t tests.

Unless otherwise stated, statistical analyses were performed in Microsoft Excel and GraphPad Prism.

